# Secretory ferritins mediate iron delivery during magnetite tooth formation in chitons

**DOI:** 10.64898/2026.10.23.761458

**Authors:** Rui Onishi, Masahiro Shimizu, Koki Okada, Nicole Angeline Robles Vizconde, Tsukasa Nakamura, Yuki Fujii, Toshiya Senda, Kenji Okoshi, Kiori Obuse, David Kisailus, Akira Satoh, Shino Homma-Takeda, Michiko Nemoto

## Abstract

Iron is essential to most organisms, yet unbound iron is toxic, necessitating tight regulation of its storage and transport. Chitons concentrate extraordinary amounts of iron in radular tissue to form magnetite-mineralized teeth, but how iron is supplied to developing teeth remains unclear. Here, synchrotron radiation X-ray fluorescence mapping resolved the spatiotemporal redistribution of iron from the radular epithelium toward developing teeth. Integrated transcriptomic and proteomic analyses identified secretory ferritin subunits in radular tissue whose genes were highly expressed in the midgut gland and whose proteins were also present in the hemolymph. In vivo fluorescence tracking showed that circulating ferritin was selectively taken up by the radular epithelium, and following injection of Ag-loaded hemolymph ferritin, Ag was detected in developing teeth. Together, these findings support a model in which secretory ferritins produced in the midgut gland transport iron through the hemolymph to the radula, enabling the pronounced iron accumulation required for tooth mineralization.

---

Chitons are marine mollusks that inhabit rocky intertidal zones and possess a feeding organ called the radula, whose teeth contain magnetite (Fig. 1a, b). These magnetite-bearing teeth exhibit greater wear resistance than zirconia and have attracted interest as models for the development of high performance structural materials ^1, 2^. As one of the few known examples of magnetite biomineralization in eukaryotes, chiton teeth provide a valuable system for investigating the biological control of iron oxide mineralization. Chitons use these hard teeth to scrape rock surfaces and feed on epilithic microalgae ^3, 4^. Teeth in the anterior region of the radula are worn away during feeding and continuously replaced by new teeth formed posteriorly. This continuous renewal results in a developmental gradient along the radula, ranging from newly formed posterior teeth to mature, functional anterior teeth ^5, 6^. Tooth formation occurs within the radular sac and is regulated by epithelial cells surrounding the developing teeth. At the posterior end of the radular sac, odontoblasts initially secrete a transparent tooth matrix composed of α-chitin and proteins. Aggregates of ferrihydrite, a poorly crystalline hydrated iron oxide, are subsequently deposited onto the chitin fibers, causing the teeth to become reddish brown. Ferrihydrite subsequently transforms into magnetite, producing mature black teeth ^7, 8^. During this process, the radular tissue accumulates more than 100 mg of iron per gram of dry tissue (>10% of dry weight) ^9, 10, 11^. Such extreme iron accumulation is exceptional among animal tissues ^12^, making the chiton radula a distinctive biological system for studying the mechanisms underlying iron-rich biomineralization.

**Fig. 1.**
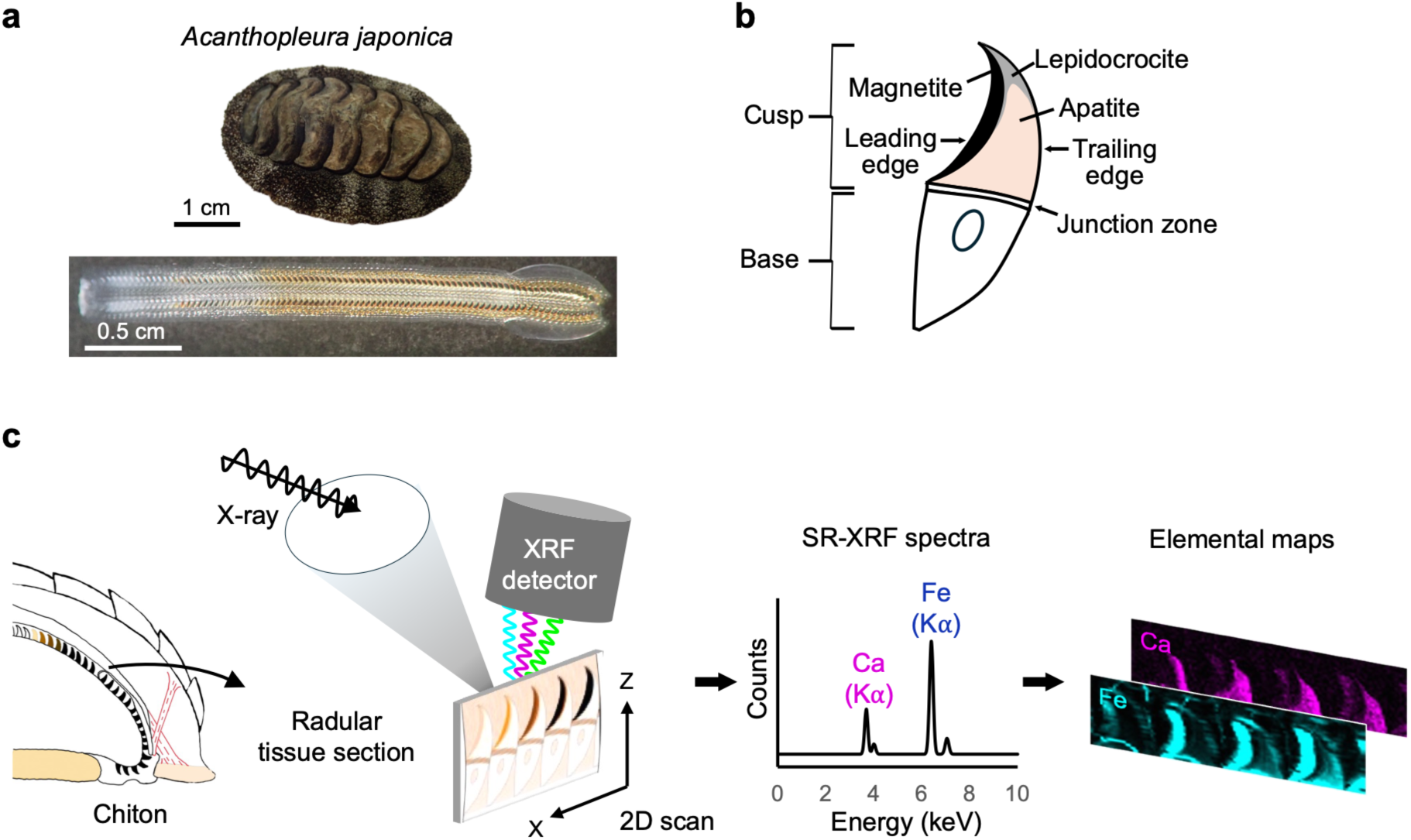
Overview of chiton radular structure and SR-XRF analysis. **a**, *Acanthopleura japonica* and its radula. **b**, Schematic representation of an *Acanthopleura*-type radular tooth. **c**, Schematic workflow for synchrotron radiation X-ray fluorescence (SR-XRF) elemental mapping of chiton radular tissue.

During tooth maturation, as iron oxide is deposited within the tooth cusps (Fig. 1b), ferritin-containing granules (siderosomes) accumulate around the developing teeth. Ferritin is a ubiquitous iron-storage protein that assembles into a multimeric nanocage structure ^13, 14, 15^. Ferritin surrounding the teeth has long been hypothesized to supply the iron required for tooth mineralization ^16, 17, 18, 19^. Mollusks possess two major ferritin types: cytoplasmic ferritin (cFer) and secretory ferritin (sFer). Cytoplasmic ferritins are localized intracellularly, whereas secretory ferritins contain signal peptides and are secreted from cells ^20^. Transcriptomic analysis of radular tissue from *Cryptochiton stelleri* revealed high expression of cytoplasmic ferritin genes ^21^. By contrast, transcriptomic analysis of radular tissue from *Acanthopleura loochooana* revealed high expression of secretory ferritin genes ^22^. Proteomic analysis of radular tissue from the same species identified both cytoplasmic and secretory ferritins ^23^. Secretory ferritins play important roles in systemic iron transport in many insects ^24, 25^ and have also been proposed to mediate iron transport in mollusks ^26, 27, 28, 29^. Despite these studies, the subunit composition of the native ferritin complexes that accumulate in radular tissue remains unknown. Moreover, whether secretory ferritins circulating in the hemolymph are taken up by the radular epithelium and contribute to iron delivery to developing teeth has not been directly demonstrated.

In the present study, we used synchrotron radiation X-ray fluorescence (SR-XRF) elemental mapping to resolve the spatiotemporal dynamics of iron within radular tissue during tooth development. By integrating biochemical isolation of native ferritin complexes with quantitative proteomics and tissue transcriptomics, we identified the secretory ferritin subunits that accumulate in radular tissue and showed that the genes encoding these subunits are highly expressed in the midgut gland. In vivo fluorescence tracking demonstrated that ferritin circulating in the hemolymph is selectively taken up by the radular epithelium. Moreover, following injection of Ag-loaded hemolymph ferritin, Ag was detected in developing teeth. Together, these findings support a model in which secretory ferritins produced in the midgut gland transport iron via the hemolymph to developing radular teeth.

## Results

### SR-XRF reveals early iron accumulation in the radular epithelium and developing teeth

To determine the spatial and temporal patterns of iron delivery during chiton tooth formation, we performed elemental mapping of the radular tissue using SR-XRF (Fig. 1c). The first tooth exhibiting visible iron oxide deposition at the leading edge was designated tooth #0; less-mature teeth were designated #−1, #−2, and #−3, etc. (with the sequence continuing toward the proximal end of the radular belt), whereas more-mature teeth were designated #1, #2, and #3, etc. (with the sequence continuing toward the distal end).

An Fe signal was first detected in the upper region of the radular epithelium overlying tooth #−16, the earliest developing tooth with a clearly recognizable tooth morphology. Fe subsequently accumulated in a dome-shaped region overlying the developing teeth. At tooth #−9, Fe accumulation first became detectable in the junction zone. Beginning at tooth #−7, a plume-like Fe distribution appeared between the junction zone and the leading edge. From tooth #−4 onward, the dome-shaped Fe-rich region shifted from the upper epithelium to epithelial cells adjacent to the teeth, while Fe deposition became detectable at the leading edge. From tooth #0 onward, Fe accumulation increased markedly in both the adjacent epithelial cells and the leading edge. Collectively, SR-XRF elemental mapping revealed that Fe begins to accumulate in radular epithelial cells and becomes detectable within developing teeth at substantially earlier developmental stages than previously recognized (Fig. 2a) ^30, 31^.

**Fig. 2.**
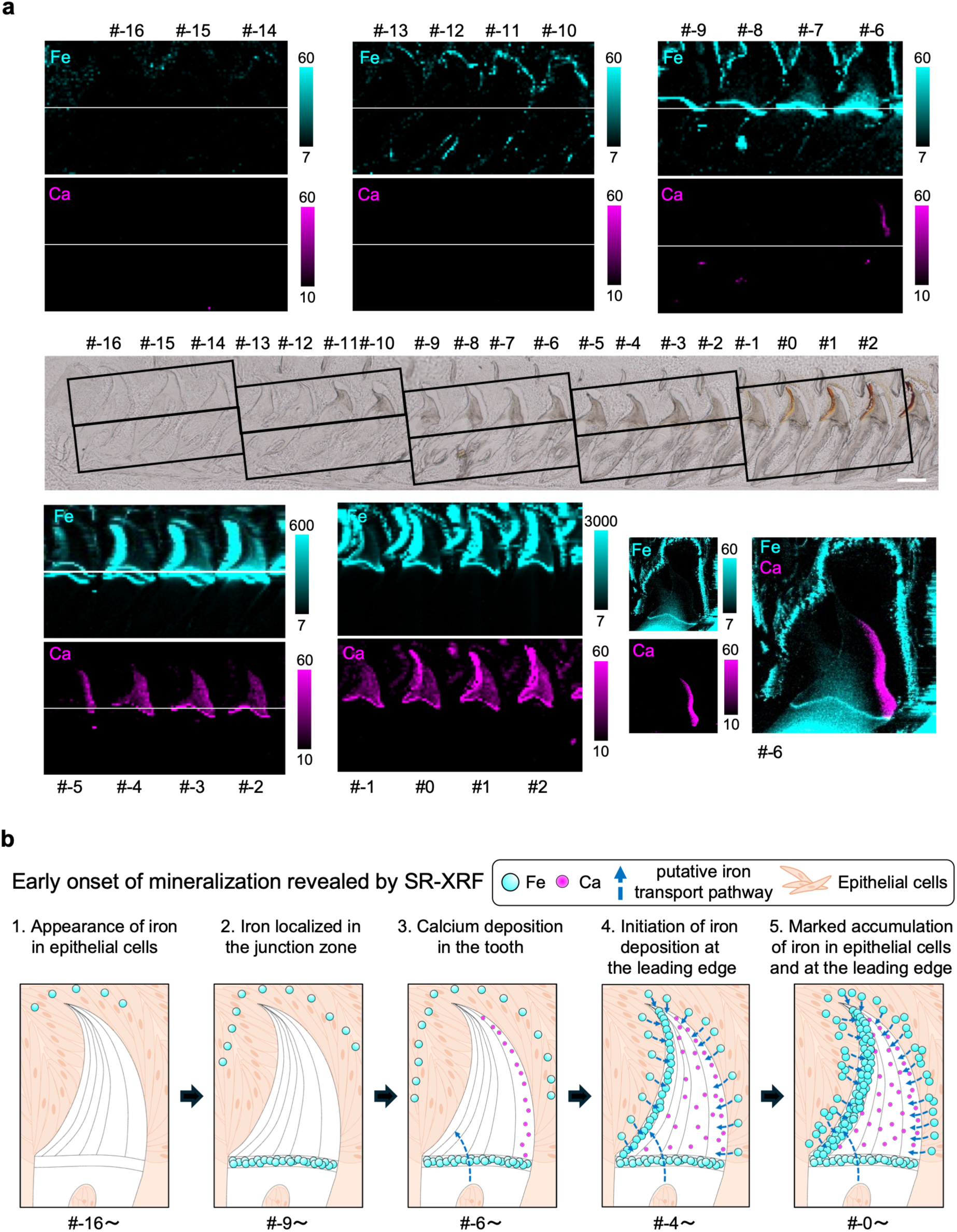
Progression of iron distribution in chiton radular tissue revealed by SR-XRF elemental mapping. **a**, X-ray fluorescence maps of Fe and Ca in chiton radular tissue. Values beside each color scale indicate the limit of quantification and the maximum X-ray fluorescence count rate for each element (counts s⁻¹). The boxed regions in the central bright-field (BF) image indicate the areas analyzed by SR-XRF. The first tooth showing visible iron oxide deposition at the leading edge by light microscopy was designated tooth #0; less-mature teeth were assigned successively lower numbers, and more-mature teeth were assigned successively higher numbers. Scale bar, 200 μm. **b**, Proposed model of Fe and Ca accumulation during radular tooth development based primarily on the SR-XRF elemental maps. The developmental progression is summarized as follows: (1) At tooth #−16, Fe first becomes detectable in the apical region of the radular epithelium. (2) At tooth #−9, Fe is distributed in a dome-shaped region overlying the tooth and first becomes detectable in the junction zone. (3) From tooth #−7 onward, a plume-like Fe distribution appears between the junction zone and the leading edge; at tooth #−6, Ca first becomes detectable on the trailing-edge side. (4) At tooth #−4, Fe first becomes detectable at the leading edge, by which stage Ca is already distributed throughout the tooth. (5) From tooth #0 onward, Fe accumulation increases markedly in both the epithelial cells adjacent to the teeth and the leading edge.

The Ca signal first became detectable on the trailing-edge side of tooth #−6 and subsequently extended into the tooth interior. At tooth #−4, when an Fe signal first became detectable at the leading edge, Ca was already distributed throughout the tooth. These observations demonstrate that Ca deposition begins earlier than previously recognized ^32^ and precedes Fe deposition at the leading edge (Fig. 2a).

The spatiotemporal progression of Fe accumulation suggested that the radular epithelium participates in iron delivery to developing teeth. Because ferritin granules in the radular epithelium surrounding developing teeth have long been proposed to supply iron for tooth mineralization ^16, 17, 18, 19^, we next sought to identify the specific ferritins present in the radular tissue.

### Identification of ferritin genes and analysis of their expression in radular tissue

To identify and classify ferritin genes in the *A. japonica* genome, we performed a homology search against its reference genome (RefSeq accession number: GCF_032854445.1) ^33^ using previously reported ferritin sequences ^21^ as queries. This search identified 10 homologous genes, each encoding a protein containing a ferritin domain of at least 150 amino acids. In the phylogenetic tree, *A. japonica* ferritins formed two distinct clades corresponding to cytoplasmic and secretory ferritins (Fig. 3a). Computational analysis identified putative iron-responsive elements (IREs) in the transcripts of one cytoplasmic and one secretory ferritin gene (Supplementary Data 2). All eight members of the secretory clade contained predicted N-terminal signal peptides and were predicted to be soluble extracellular proteins (Supplementary Data 3). Sequence alignment identified a characteristic insertion in the secretory ferritins, which AlphaFold 3 modeling predicted to form a non-helical loop (Fig. 3b). Together, these analyses indicated that the *A. japonica* genome encodes two cytoplasmic and eight secretory ferritins.

**Fig. 3.**
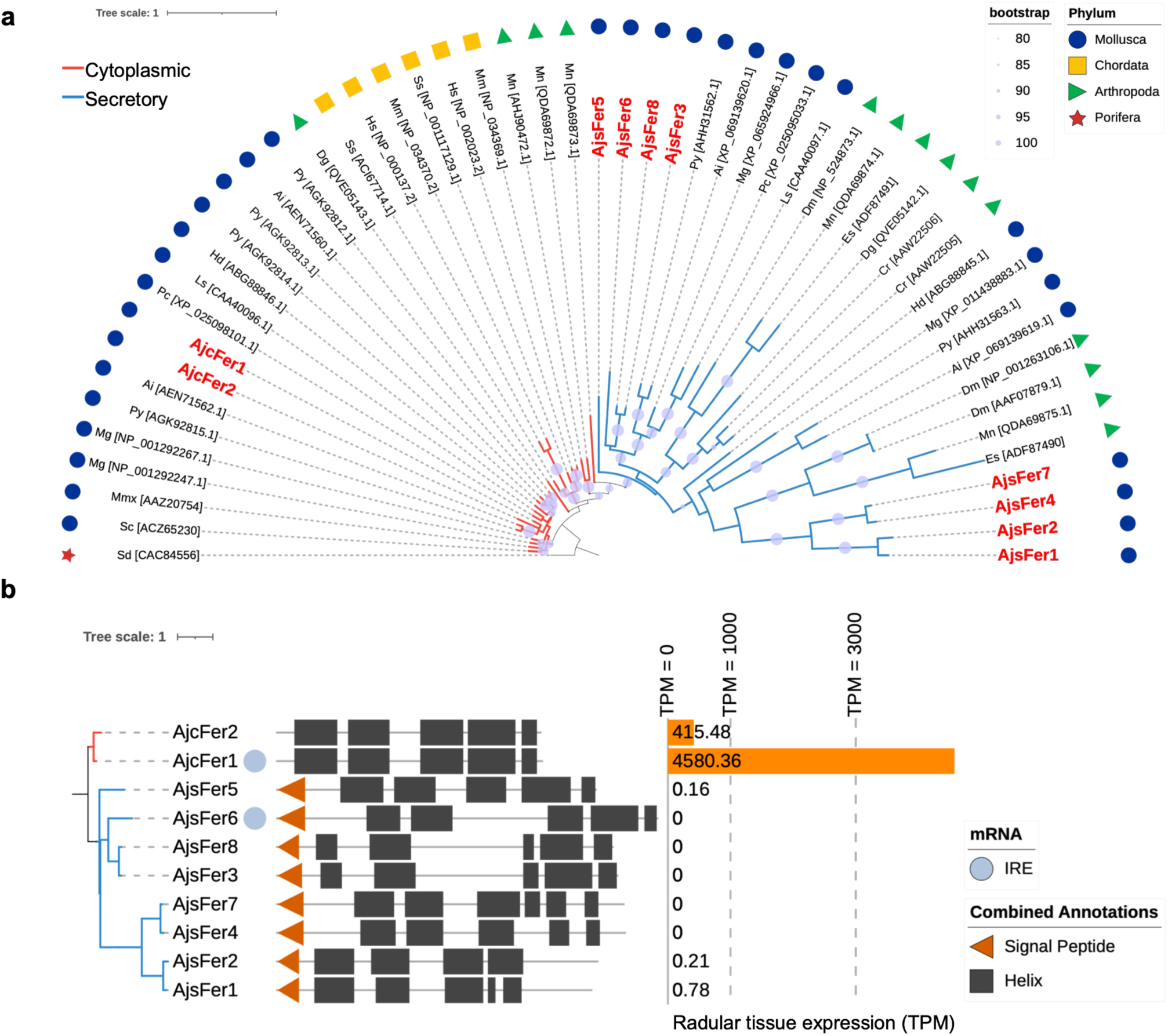
Phylogenetic analysis and classification of ferritin genes in *A. japonica* and expression analysis in radular tissue. **a**, Phylogenetic tree constructed from 53 ferritin amino acid sequences representing *A. japonica* and other molluscs, arthropods, vertebrates, and a poriferan outgroup. The tree resolved two major clades corresponding to cytoplasmic ferritins (red branches) and secretory ferritins (blue branches). Branch support values are shown as percentages. Phyla are indicated by symbols and colors, and *A. japonica* ferritins are highlighted in red. The sequences used for phylogenetic analysis are listed in Supplementary Data 1. **b**, Classification and sequence-based structural annotation of the *A. japonica* ferritins, together with the presence or absence of predicted iron-responsive elements (IREs). The tree on the left was pruned from the tree shown in a using iTOL to display only the *A. japonica* ferritins. The bar graph on the right shows transcript abundance in radular tissue, expressed as transcripts per million (TPM), calculated by mapping quality-filtered RNA-seq reads to the *A. japonica* reference genome. cFer, cytoplasmic ferritin; sFer, secretory ferritin.

We next analyzed RNA-seq data from *A. japonica* radular tissue to characterize the expression levels of the identified ferritin genes ^7^. After quality filtering, the reads were mapped to the reference genome, and transcript abundance was quantified in transcripts per million (TPM). Of the two cytoplasmic ferritin genes, one was highly expressed in radular tissue, whereas the eight genes encoding secretory ferritins showed little to no expression (Fig. 3b). A similar expression pattern was observed in all four chiton species examined (Supplementary Fig. 1).

### Isolation and characterization of native ferritin complexes from radular tissue

Following the analysis of ferritin gene expression, we isolated native ferritin complexes from radular tissue and characterized their biochemical properties and subunit composition. Epithelial cells isolated from radular tissue were disrupted by ultrasonication and subjected to heat treatment. After centrifugation, the supernatant was ultracentrifuged at 100,000 × *g*, and the resulting pellet was resuspended in PBS and subjected to size-exclusion chromatography. Size-exclusion chromatography revealed a peak at an elution volume corresponding to an apparent molecular mass of ≤ 670 kDa (Supplementary Fig. 2a). Blue native polyacrylamide gel electrophoresis (BN-PAGE) showed that the native radular ferritin complex had a higher apparent molecular mass than horse spleen ferritin (480 kDa) (Fig. 4a). Prussian blue staining following native PAGE confirmed that the radular ferritin complex contained iron (Fig. 4b). SDS-PAGE revealed a single major band at approximately 25 kDa, consistent with the molecular mass of a ferritin subunit (Fig. 4c). TEM revealed spherical structures with cores 7−10 nm in diameter (Fig. 4d).

**Fig. 4.**
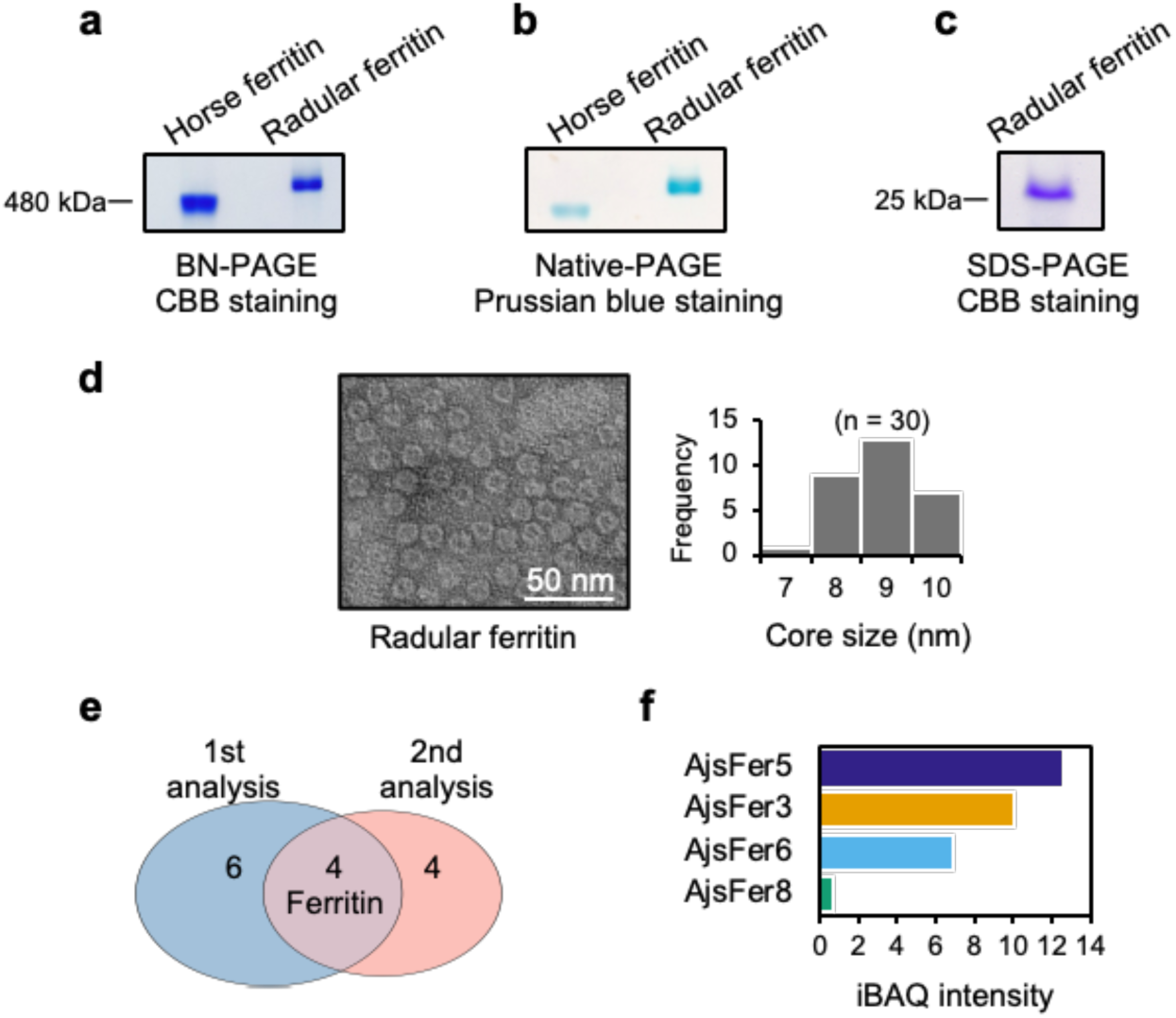
Isolation and proteomic characterization of native ferritin complexes from radular tissue. **a**, Blue native PAGE (BN-PAGE) followed by Coomassie Brilliant Blue (CBB) staining. **b**, Native PAGE followed by Prussian blue staining. Horse spleen ferritin was included as a reference in **a** and **b**. **c**, Tricine SDS-PAGE followed by CBB staining. **d**, Representative transmission electron microscopy (TEM) image of ferritin isolated from radular tissue and the corresponding distribution of ferritin core diameters. A total of 30 particles were measured (n = 30). Scale bar, 50 nm. **e**, Venn diagram showing the proteins identified by nanoLC–MS/MS in two independently prepared biological replicates. The approximately 25-kDa band separated by Tricine SDS-PAGE was analyzed, and four ferritin subunits were identified in both replicates. **f**, Intensity-based absolute quantification (iBAQ) values of the four ferritin subunits identified in both replicates. Values shown were calculated from the MS data obtained for replicate 1.

We next used N-terminal sequencing and nanoLC-MS/MS to determine the subunit composition of the purified ferritin complexes. N-terminal sequencing of the approximately 25-kDa band separated by SDS-PAGE (Fig. 4c) yielded the sequence EDKRL, which matched the predicted mature N terminus of AjsFer3. NanoLC–MS/MS analysis was performed using two independently prepared biological replicates. In both replicates, all ferritin-derived peptides were assigned to secretory ferritins (Fig. 4e). Moreover, the rank order of the Intensity-Based Absolute Quantification (iBAQ) values for the identified secretory ferritins was identical between the replicates (Supplementary Data 5). These results demonstrate that native ferritin complexes isolated from radular tissue are composed predominantly of secretory ferritin subunits (Fig. 4f).

### Tissue expression profiles of secretory ferritin genes

The genes encoding the secretory ferritin subunits identified in native radular ferritin complexes showed little to no expression in radular tissue (Fig. 3b). We therefore used RT-qPCR to examine the tissue distribution of transcripts encoding AjsFer3, one of these subunits. Four tissues were analyzed: radular tissue, midgut gland, gonad, and foot (Fig. 5a). The abundance of transcripts encoding AjsFer3 was highest in the midgut gland (Fig. 5b). We then examined the tissue expression profiles of the remaining secretory ferritin genes by performing RNA-seq analysis of the midgut gland and foot and comparing these data with those obtained from radular tissue. Quality-filtered reads were mapped to the *A. japonica* reference genome (RefSeq accession number: GCF_032854445.1) ^33^, using the same analytical pipeline as that used for radular tissue. Both cytoplasmic ferritin genes were expressed in all three tissues, with the IRE-containing *AjcFer1* showing higher expression. By contrast, the secretory ferritin genes, which showed little to no expression in radular tissue, were highly expressed in the midgut gland and foot (Fig. 5c). Notably, all genes encoding the secretory ferritin subunits detected in native radular ferritin complexes were highly expressed in the midgut gland. These expression profiles suggest that the secretory ferritins detected in radular tissue may originate in the midgut gland and be transported to the radula via the hemolymph.

**Fig. 5.**
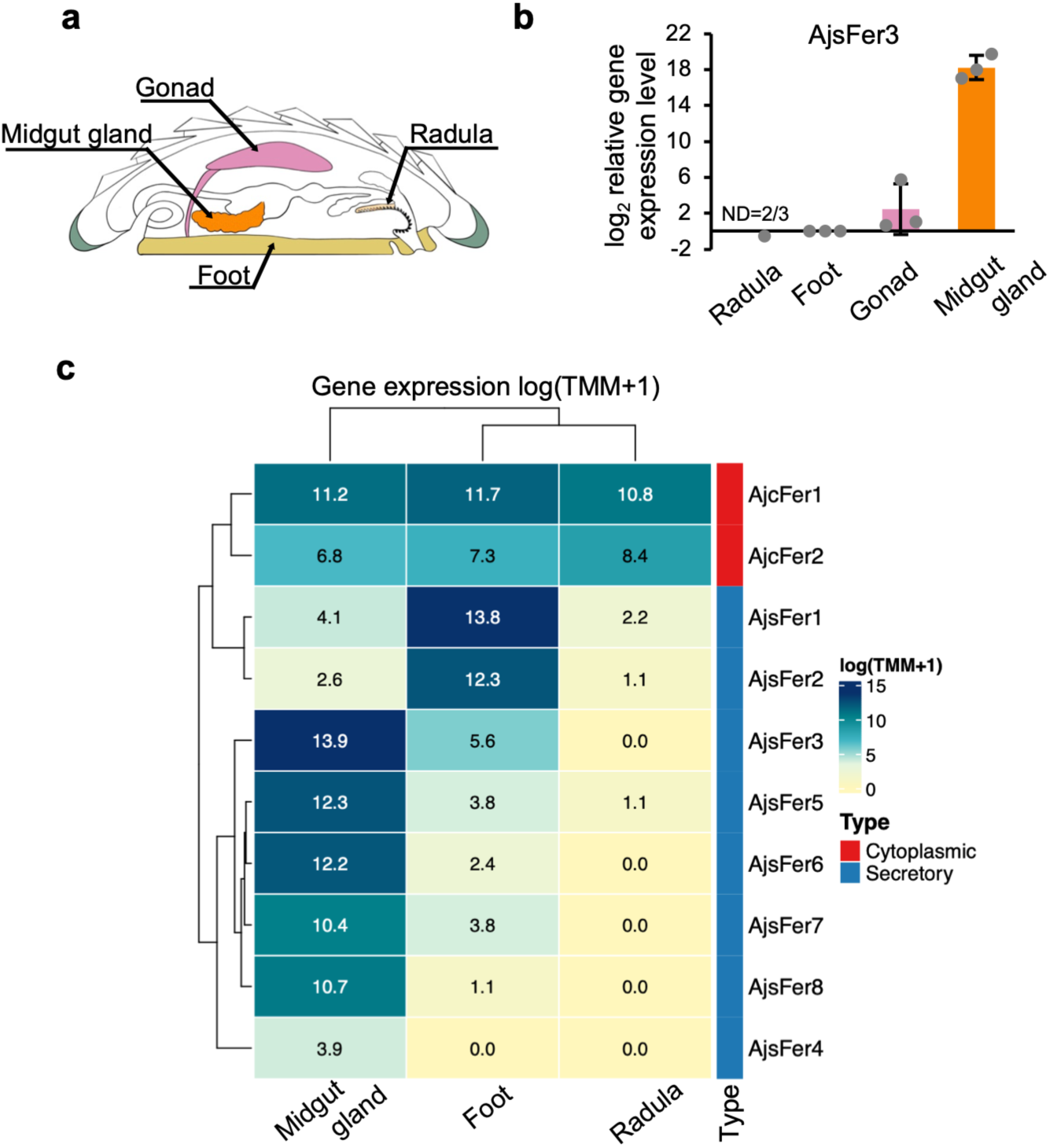
Tissue expression profiles of *A. japonica* ferritin genes. **a**, Cross-sectional schematic of *A. japonica* showing the tissues analyzed: midgut gland, gonad, radula, and foot. **b**, RT-qPCR analysis of transcripts encoding AjsFer3 in the indicated tissues. Transcript abundance was normalized to that of the actin reference gene. Data are presented as mean ± SD (n = 3 biologically independent individuals). ND, not detected. **c**, Heatmap showing ferritin gene expression in the midgut gland, foot, and radular tissue. Colors represent natural log-transformed TMM-normalized counts per million [log (TMM-normalized CPM + 1)]. The dendrogram on the left represents hierarchical clustering based on similarities in expression profiles and does not indicate phylogenetic relationships. Row annotations indicate cytoplasmic ferritins in red and secretory ferritins in blue.

### Isolation and characterization of native ferritin complexes from hemolymph

We next tested this hypothesis by examining whether the secretory ferritins detected in radular tissue were also present in the hemolymph. Native ferritin complexes were purified from the hemolymph using the same procedure as that used for radular ferritin, except that the ultrasonication step was omitted. Size-exclusion chromatography revealed a peak at an elution volume similar to that of the radular ferritin peak (Supplementary Fig. 2e). BN-PAGE showed that hemolymph ferritin formed a complex with an apparent molecular mass greater than 480 kDa (Fig. 6a), similar to radular ferritin. Prussian blue staining following native PAGE likewise confirmed that the hemolymph ferritin complex contained iron (Fig. 6b). In contrast to the single major band observed for radular ferritin, SDS-PAGE of hemolymph ferritin revealed three bands at approximately 25 kDa (Fig. 6c). TEM revealed spherical structures with cores 7-10 nm in diameter, comparable to those of radular ferritin (Fig. 6d).

**Fig. 6.**
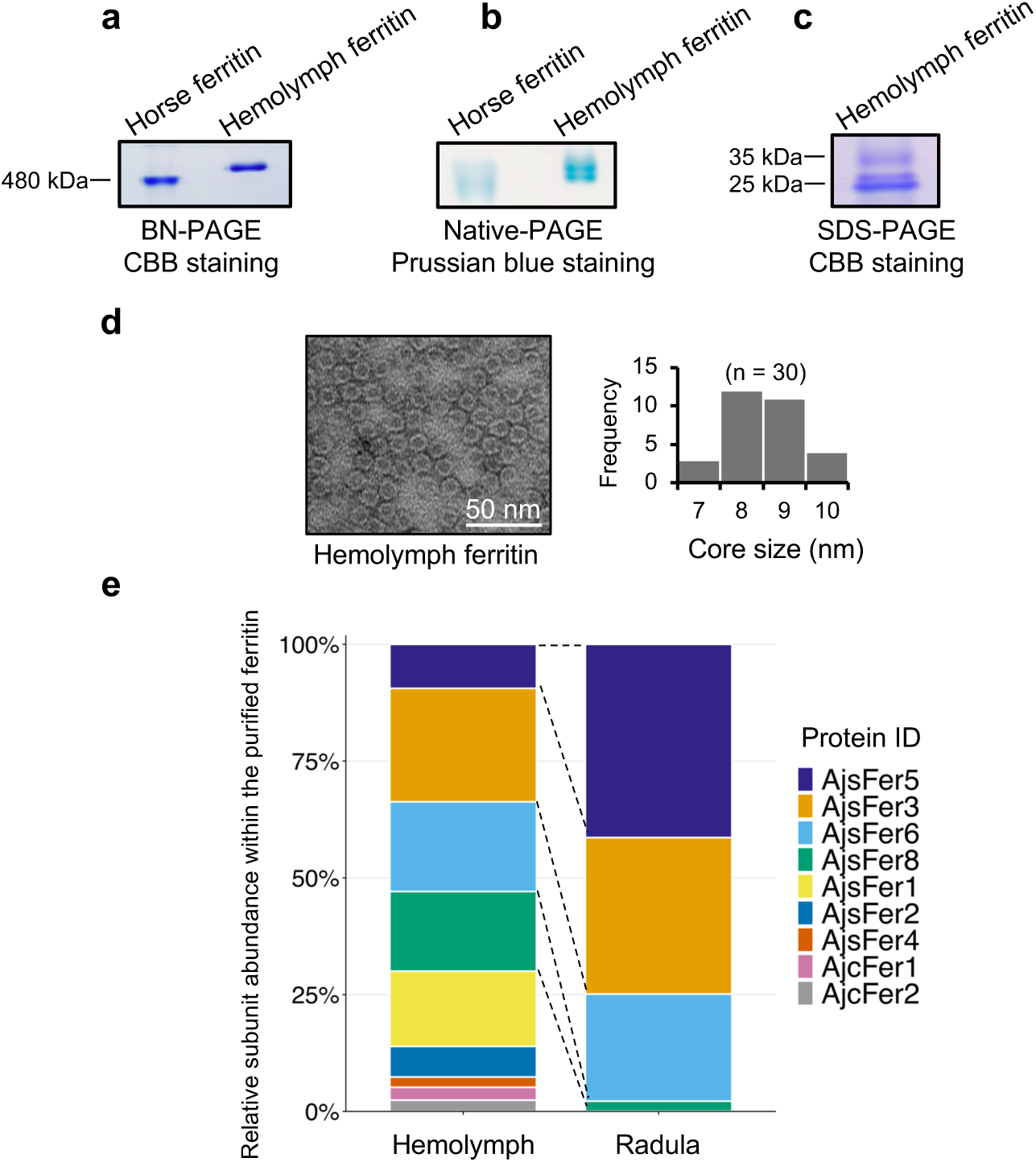
Isolation and proteomic characterization of native ferritin complexes from hemolymph. **a**, BN-PAGE followed by CBB staining. **b,** Native PAGE followed by Prussian blue staining. **c**, Tricine SDS-PAGE followed by CBB staining. **d**, Representative TEM image of ferritin isolated from hemolymph and the corresponding distribution of ferritin core diameters. A total of 30 particles were measured (n = 30). Scale bar, 50 nm. **e**, Relative ferritin subunit compositions of the purified hemolymph and radular ferritin preparations. Relative iBAQ (riBAQ) values were calculated by dividing the iBAQ value of each ferritin subunit by the summed iBAQ values of all ferritin subunits detected in the corresponding proteomic dataset. Dashed lines connect the four subunits detected in both hemolymph and radular ferritin preparations.

Because overlapping sequence signals prevented the identification of individual subunits by N-terminal sequencing, the three bands detected by SDS-PAGE were analyzed separately by nanoLC– MS/MS using the same procedure as that used for radular ferritin. Two cytoplasmic and seven secretory ferritin subunits were identified in the hemolymph ferritin complexes. The genes encoding all seven secretory ferritins were expressed in the midgut gland (Fig. 5c). All four secretory ferritin subunits identified in radular ferritin complexes were also detected in hemolymph ferritin complexes (Fig. 6e). In the phylogenetic tree, these four shared subunits clustered together within the secretory ferritin clade (Fig. 3a). These results support a model in which a specific subset of secretory ferritins circulating in the hemolymph is preferentially transported to radular tissue.

### Identification of candidate motifs associated with ferritin transport to radular tissue

To identify sequence features potentially associated with selective transport to radular tissue, we compared the seven secretory ferritin subunits detected in the hemolymph. These subunits were divided into two groups: four that were also detected in radular tissue and three that were detected only in the hemolymph. Motif analysis identified three motifs present in the radula-associated subunits but absent from the hemolymph-only subunits (Fig. 7a, b).

**Fig. 7.**
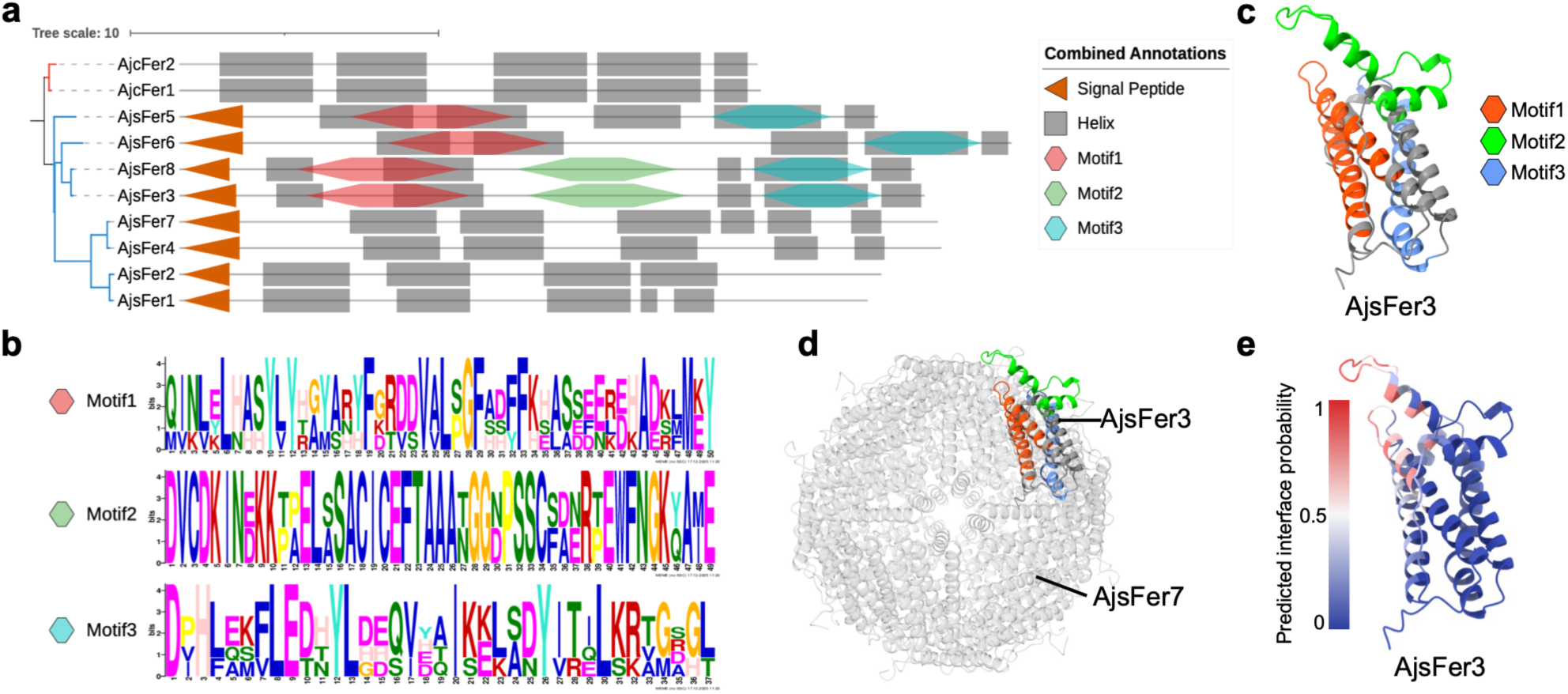
Structural characterization of radula-associated ferritins in *A. japonica*. **a**, Phylogenetic relationships and sequence-based structural annotations of the ten *A. japonica* ferritins. Orange triangles indicate predicted signal peptides, gray boxes indicate predicted α-helical regions, and colored regions indicate the three candidate radula-associated motifs identified using MEME and mapped using FIMO. The motif sequences and associated *P* values are provided in Supplementary Data 8. **b**, Sequence logos of the three candidate motifs. The total height of each letter stack represents the information content at that position, and the relative height of each amino acid letter reflects its frequency. **c**, AlphaFold 3-predicted structural model of AjsFer3 showing the locations of motifs 1, 2, and 3 in red, green, and blue, respectively. **d**, Cryo-electron microscopy structure of the recombinant AjsFer7 24-mer, with the AlphaFold 3 model of AjsFer3 superimposed onto one AjsFer7 subunit. **e**, Per-residue probabilities of involvement in protein– protein interaction interfaces predicted using PeSTo, with red indicating higher and blue indicating lower predicted probabilities.

We next mapped these motifs onto an AlphaFold 3 model of AjsFer3, one of the radula-associated ferritin subunits (Fig. 7c). Motifs 1 and 3 were located within the four-helix bundle, whereas motif 2 was located in an extended non-helical loop between helices B and C (the B−C loop). Motif 2 was characterized by the presence of four cysteine residues. To assess the position of motif 2 within the assembled ferritin complex, we superimposed the AlphaFold 3 model of AjsFer3 onto the cryo-EM structure of recombinant AjsFer7, determined at an overall resolution of 2.00 Å (Supplementary Fig. 3 and Supplementary Table 1). Although the mature AjsFer3 and AjsFer7 sequences shared only 21.7% amino acid sequence identity across 263 aligned residues, their four-helix cores closely superimposed. Structural deviations were concentrated in non-helical regions, and the root-mean-square deviation (RMSD) was 0.885 Å after exclusion of the variable non-helical regions. The conserved core structure supported placement of the AjsFer3 model within the AjsFer7 24-mer assembly, in which motif 2 was predicted to be positioned on the outer surface of the ferritin cage (Fig. 7d). Finally, PeSTo analysis ^34^ identified several residues within motifs 1 and 2 as potential protein-protein interaction sites (Fig. 7e). These analyses identified candidate motifs that may mediate the selective recognition of circulating ferritins by a putative receptor and their subsequent uptake into radular tissue.

### In vivo tracking of hemolymph ferritin transport and Ag delivery

We tracked the distribution of Alexa Fluor 488-labeled hemolymph ferritin after injection into *A. japonica* to test whether it is transported to radular tissue in vivo (Fig. 8a). Alexa Fluor 488-labeled horse spleen ferritin served as a control. At 48 h after injection, fluorescence from labeled hemolymph ferritin was detected around developing teeth #0–#4 and closely coincided with the ferritin-rich granular regions visible in bright-field images (Fig. 8b, Supplementary Figs. 4 and 5). By contrast, labeled horse spleen ferritin did not accumulate around the teeth. Instead, its fluorescence was restricted to the upper region of the radular epithelium overlying teeth #−3 to #−1. This distribution may reflect nonspecific adsorption to the hemolymph-facing surface of the epithelium. These results indicate that hemolymph ferritin is selectively transported to radular tissue and accumulates in ferritin-rich regions surrounding developing teeth.

**Fig. 8.**
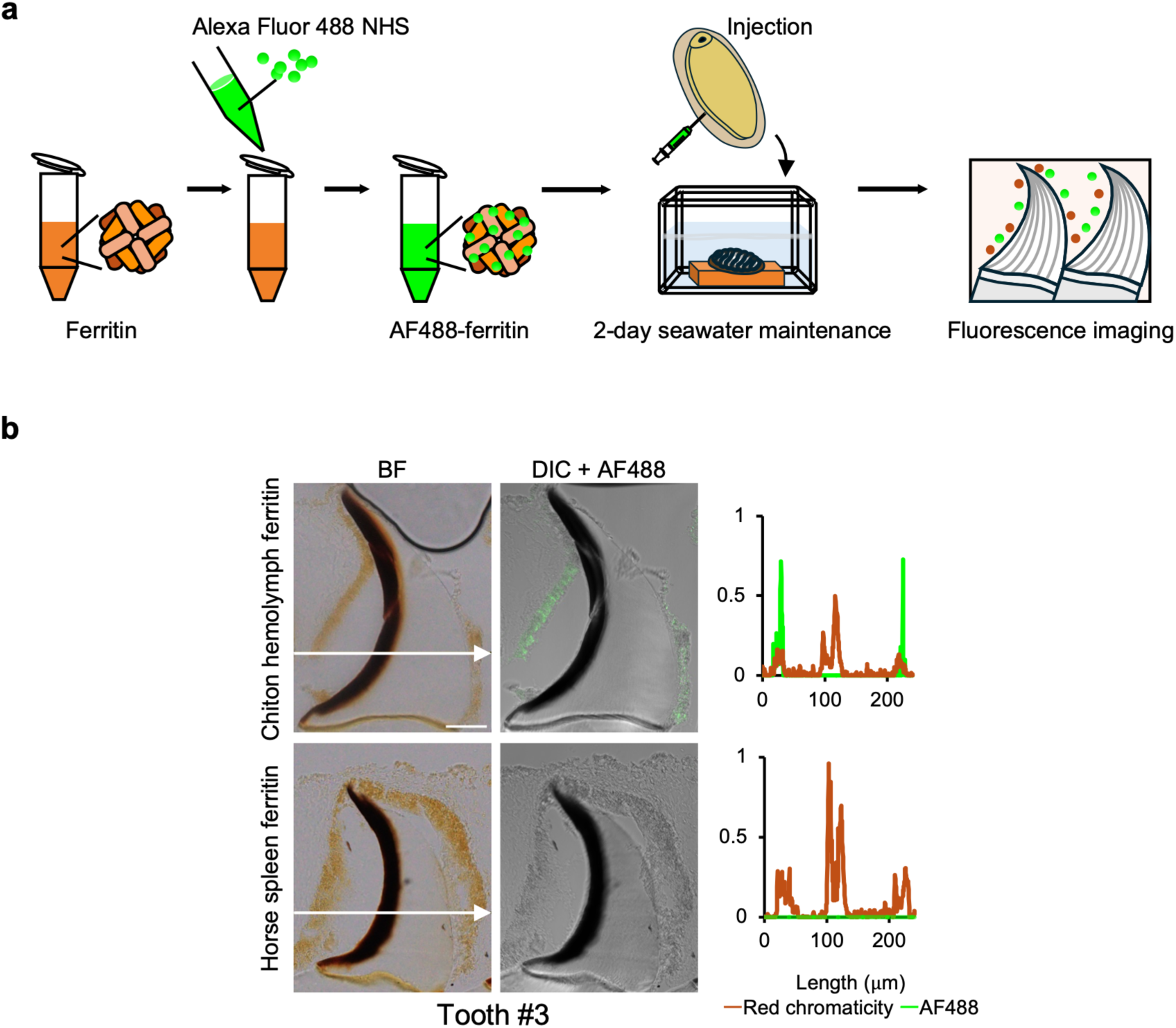
In vivo tracking of fluorescently labeled hemolymph ferritin in radular tissue. **a**, Schematic representation of the in vivo ferritin-tracking experiment. **b**, Representative BF and differential interference contrast (DIC)/Alexa Fluor 488 images of tooth #3 from chitons injected with fluorescently labeled chiton hemolymph ferritin or horse spleen ferritin. Images of all examined teeth are shown in Supplementary Figs. 4 and 5. The graphs on the right show normalized spatial profiles of Alexa Fluor 488 fluorescence (green) and red chromaticity (red), measured along the arrows shown in the BF images. Red chromaticity indicates the ferritin-rich granular regions surrounding the tooth and the iron oxide-rich region within the tooth. Scale bar, 50 μm.

To investigate whether hemolymph ferritin contributes to iron delivery to developing teeth, we loaded purified hemolymph ferritin with Ag as an elemental tracer and injected it into the hemolymph (Fig. 9a). Iron removal and Ag loading were confirmed by ICP-MS (Supplementary Table 2), and TEM showed that the ferritin complexes retained their overall morphology after Ag loading (Supplementary Fig. 6). SR-XRF elemental mapping performed 3 days after injection revealed no distinct Ag signal in tooth #−3, whereas Ag was detected at the leading edge and junction zone of tooth #0 (Fig. 9b). In the apical region of tooth #0, Ag was also detected on the trailing-edge side, where no Fe signal was observed. In the more mature teeth #1 and #2, the Ag signal at the leading edge became progressively more pronounced. In tooth #2, Ag was also detected on the trailing-edge side, in addition to the leading edge and junction zone. Ag localization in tooth #0 was consistently observed in three independent individuals (Supplementary Fig. 7a). Spectral analysis confirmed that Ag was detected in individuals injected with Ag-loaded hemolymph ferritin but not in the uninjected control (Supplementary Fig. 7b). These results show that Ag initially loaded into hemolymph ferritin reached developing teeth, supporting the hypothesis that hemolymph ferritin contributes to iron delivery during tooth mineralization.

**Fig. 9.**
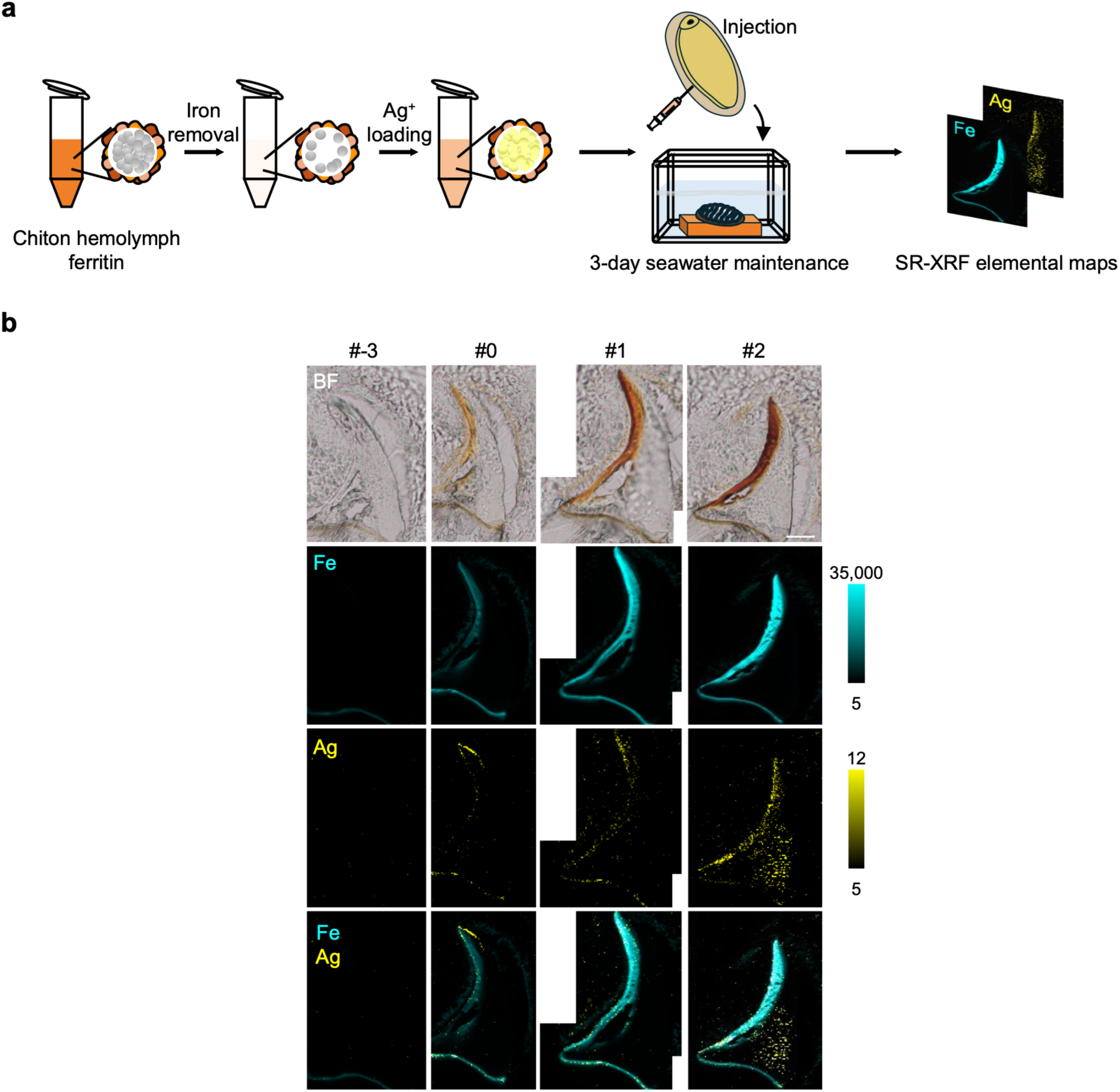
SR-XRF mapping of Fe and Ag in radular teeth following injection of Ag-loaded hemolymph ferritin. **a**, Schematic representation of ferritin deironing, Ag loading, injection, and subsequent SR-XRF elemental mapping. **b**, BF images, X-ray fluorescence maps of Fe and Ag, and merged Fe/Ag maps of teeth #−3, #0, #1, and #2. Values beside each color scale indicate the limit of quantification and the maximum X-ray fluorescence count rate (counts s⁻¹). Scale bar, 50 μm.

## Discussion

SR-XRF analysis revealed that iron accumulation in radular epithelial cells begins at a substantially earlier stage of tooth development than previously recognized. In a previous TEM study of radular tissue from *Acanthopleura hirtosa*, ferritin granules were first observed at a developmental stage approximately corresponding to tooth #−6 in the numbering scheme used in this study ^17^. By contrast, SR-XRF mapping detected Fe in the radular epithelium overlying tooth #−16, the earliest developing tooth with a clearly recognizable morphology in our analysis (Fig. 2a). At this stage, the Fe signal appeared as discrete spots in the apical region of the epithelium. As tooth development advanced from tooth #−16 to #−10, these spots increased in number and shifted progressively toward the developing teeth. To our knowledge, these maps provide the first detailed visualization of the stage-dependent shift in iron distribution from the apical region of the radular epithelium toward developing teeth.

Iron delivered to developing teeth is thought to accumulate first in the junction zone, which serves as a temporary reservoir, before being supplied to the cusp region. The Fe accumulated in this region has not been assigned to a distinct mineral phase by Raman spectroscopy ^30, 31, 35, 36^. Earlier work detected Fe in the junction zone of tooth #−5 but observed a plume-like Fe distribution connecting the junction zone and leading edge only at tooth #0, coincident with the initial appearance of iron mineral at the leading edge ^30^. By contrast, our SR-XRF maps detected Fe in the junction zone as early as tooth #−9 and resolved a plume-like Fe distribution at tooth #−7, before an Fe signal first became detectable at the leading edge of tooth #−4 (Fig. 2a). These observations resolve iron delivery to developing teeth into three temporally distinct stages: accumulation in the junction zone, formation of a plume-like distribution, and accumulation at the leading edge, with the plume-like distribution representing an intermediate stage in this sequence.

Previous studies have suggested that Ca is incorporated into the tooth core after the onset of Fe deposition at the leading edge ^30, 31, 35, 36^. In our SR-XRF maps, however, Ca was already detectable on the trailing-edge side of tooth #−6, a stage at which a plume-like Fe distribution was present but leading-edge Fe deposition had not yet begun (Fig. 2a). These observations indicate that Ca accumulation begins earlier than previously recognized, during an early phase of Fe delivery. The spatially distinct distributions of Fe and Ca in tooth #−6 further suggest that the transport and deposition of these elements are differentially regulated within developing teeth.

RNA-seq analysis showed that the predominant ferritin transcripts in *A. japonica* radular tissue were derived from cytoplasmic ferritin genes (Fig. 3b). This finding is consistent with transcriptomic data from the radular tissue of another chiton species, *Cryptochiton stelleri* ^21^. For each of the two cytoplasmic ferritin genes, transcript abundance was broadly similar among the midgut gland, foot, and radular tissue. Thus, the high abundance of cytoplasmic ferritin transcripts in radular tissue likely reflects basal expression maintained across tissues rather than radula-specific upregulation. In all three tissues, transcripts of the IRE-containing gene *AjcFer1* were more abundant than those of the IRE-lacking gene *AjcFer2*. Similar expression patterns have been reported for cytoplasmic ferritins in *Acanthopleura granulata* ^37^. Despite the abundance of cytoplasmic ferritin transcripts in the radula, cytoplasmic ferritin subunits were not detected in native ferritin complexes isolated from radular tissue; instead, these complexes were composed predominantly of secretory ferritin subunits. One possibility is that sequestration of iron within secretory ferritin-containing granules limits cytosolic iron availability and thereby suppresses the translation of IRE-containing *AjcFer1*, although this hypothesis requires direct experimental testing.

Chitons, including *A. japonica*, inhabit rocky substrates and feed primarily on epilithic microalgae ^10^. The iron required for tooth formation is thought to derive from several sources, including ingested algae, rock particles abraded during grazing, seawater, and iron recycled from worn teeth ^11^. The molluscan midgut gland plays central roles in digestion, nutrient absorption, and nutrient storage ^38, 39, 40,41^. Radiotracer studies in other mollusks have also demonstrated iron uptake by digestive-tract epithelial cells followed by transfer to the hemolymph ^42^. These observations suggest that the midgut gland is a likely site of iron uptake in chitons, although the chemical form and cellular mechanism of iron uptake remain unknown.

In *A. japonica*, the genes encoding the secretory ferritin subunits detected in radular tissue were highly expressed in the midgut gland, and the same subunits were detected in the hemolymph. These findings suggest that iron absorbed through the midgut gland is incorporated into secretory ferritin and subsequently released into the hemolymph. Previous reports of high ferritin concentrations in chiton hemolymph further support a physiological role for ferritin as a circulating iron carrier ^18, 43^. A similar systemic iron transport mechanism is well established in insects, in which ferritin assembles within the secretory pathway, incorporates iron absorbed by the midgut, and is secreted into the hemolymph for delivery to other tissues ^24, 25, 44, 45, 46, 47^. Related ferritin transport pathways have also been described in gastropods, where yolk ferritin synthesized in the midgut gland is secreted into the hemolymph and subsequently taken up by oocytes ^29^. Our findings extend this concept to chiton tooth mineralization by providing direct in vivo evidence that ferritin circulating in the hemolymph is selectively taken up by radular tissue (Fig. 8, 9).

Proteomic analysis of native ferritin complexes isolated from radular tissue identified four secretory ferritin subunits. Ferritin generally assembles into a 24-mer cage, and X-ray crystallographic analyses of recombinant molluscan ferritins have shown that this fundamental structure is conserved ^48, 49, 50^. Mammalian ferritins often form heteropolymeric cages whose subunit compositions vary among tissues ^14^. The four subunits detected in *A. japonica* radular ferritin may therefore co-assemble into heteromeric 24-mers. The reproducible relative iBAQ profiles may reflect the average subunit composition of the purified ferritin population but do not directly establish the number or proportion of each subunit within individual 24-mers. Further structural and biochemical analyses of native radular ferritin will be required to resolve its subunit organization. The cryo-EM reconstruction of recombinant AjsFer7 reported here provides a structural framework for future analyses of chiton ferritins.

Sequence analysis identified three motifs that were present in the radula-associated ferritin subunits but absent from the hemolymph-only subunits. Motifs 1 and 2 contain amino acid residues predicted to participate in protein-protein interaction interfaces, and motif 2 is located in the B−C loop of the ferritin structure. In mammals, transferrin receptor 1 (TfR1) functions as a receptor for H-ferritin, and structural analysis has shown that the B−C loop contributes to the ferritin–TfR1 binding interface ^51^. This positional correspondence raises the possibility that motif 2 contributes to the recognition of circulating ferritin by a putative receptor on radular epithelial cells. Previous TEM observations of chiton radular epithelial cells have suggested that ferritin is internalized by endocytosis ^52^. Moreover, only a subset of the secretory ferritin subunits circulating in the hemolymph was detected in radular tissue in the present study, consistent with selective ferritin uptake. A previous study similarly found that chiton hemolymph contains multiple ferritin types, whereas particular ferritins show distinct tissue distributions ^53^. Together, these findings support a model in which secretory ferritins originating from the midgut gland circulate in the hemolymph, are selectively internalized by radular epithelial cells, potentially through receptor-mediated endocytosis, and contribute to iron delivery to developing teeth (Supplementary Fig. 8). This pathway may help account for the pronounced iron enrichment of chiton radular tissue ^9, 10, 53^. Although candidate ferritin receptors expressed in molluscan hemocytes have been reported, whether they mediate ferritin uptake in other tissues remains unknown ^27^. The motifs identified in this study represent candidate recognition elements for future studies aimed at identifying the radular ferritin receptor and elucidating the mechanism of tissue-specific ferritin uptake.

Fluorescence derived from labeled hemolymph ferritin was detected in radular epithelial cells but not within the developing teeth. Previous TEM observations of radular tissue in *A. hirtosa* similarly showed that ferritin particles were associated with microvilli surrounding sites of magnetite deposition but were absent from the tooth interior ^17, 54^. Proteomic analyses of tooth cusps likewise failed to detect ferritin ^21, 55^. Cryo-TEM observations of the limpet *Patella caerulea* further revealed particles consistent in size with ferritin cores in tissue adjacent to the teeth but no such particles within the tooth matrix ^56^. Moreover, the iron initially deposited in radular teeth has been characterized as a low-nuclearity complex rather than the ferrihydrite-like mineral characteristic of ferritin cores ^57^. Together, our fluorescence-tracking results and previous ultrastructural, proteomic, and chemical evidence support a model in which iron is released from ferritin before being incorporated into the developing tooth, whereas the intact ferritin complex does not enter the tooth matrix.

Following injection of Ag-loaded hemolymph ferritin, SR-XRF mapping detected Ag in developing teeth (Fig. 9b), supporting the involvement of hemolymph ferritin in iron delivery during tooth mineralization. Although the Ag signal at the leading edge increased with tooth maturation, its distribution remained broader than that of Fe and lacked the strong leading-edge enrichment characteristic of Fe. This difference may reflect metal selectivity in the downstream mineralization process. Because RTMP1 binds Fe^3+^ and promotes iron oxide nucleation at the leading edge ^7^, it may interact less efficiently with Ag-containing species than with Fe^3+^, although this possibility requires direct experimental testing.

For more than half a century, ferritin has been proposed to supply iron for chiton tooth mineralization, but the specific ferritins involved had not been identified ^19^. In the present study, we identified the secretory ferritin subunits present in radular tissue and provided direct in vivo evidence that ferritin circulating in the hemolymph is selectively taken up by the radular epithelium. These findings support a ferritin-mediated pathway that transports iron through the hemolymph to the radula for tooth mineralization, providing new insight into systemic iron transport and tissue-specific iron accumulation in animals.

## Methods

### SR-XRF Analysis of Chiton Radular Tissue

Specimens of *A. japonica* were collected from the intertidal zone around Ushimado Harbor, Okayama Prefecture, Japan. All experiments involving chitons were conducted in accordance with institutional guidelines and were approved by the Animal Care and Use Committee of Okayama University (approval no. OKU-2023651).

Freshly dissected radular tissues were immediately fixed in 4% paraformaldehyde in phosphate-buffered saline (PBS) at 4°C for 6 h. The tissues were then dehydrated through a graded ethanol series to 100% ethanol and embedded in paraffin. Four-micrometer-thick sections were prepared from the paraffin-embedded tissues, mounted on polypropylene film, and air-dried. The sections were then subjected to SR-XRF elemental mapping.

SR-XRF measurements were performed at beamline BL37XU of SPring-8 (Japan Synchrotron Radiation Research Institute, Hyogo, Japan). The incident X-rays were monochromatized to 37 keV using a Si (111) double-crystal monochromator and focused to a beam size of 1 × 1 μm at the sample position using Kirkpatrick–Baez mirrors. The incident X-ray intensity was monitored using an ionization chamber. For the large-area elemental maps shown in Fig. 2a, the tissue sections were scanned with a step size of 10 μm in both the X and Z directions. For the enlarged map of tooth #−6 shown in Fig. 2a and the analyses shown in Fig. 9b and Supplementary Fig. 7a, the sections were scanned with step sizes of 2 μm in both the X and Z directions. At each measurement point, an X-ray fluorescence spectrum was acquired for 1 s using a silicon drift detector (SDD). Elemental maps were generated from the Fe Kα, Ca Kα, and Ag Kα lines measured at each point. Fiji ^58^ was used to generate the elemental maps. Calibration curves for Fe, Ca, and Ag were constructed from their respective Kα lines obtained by scanning standard samples containing 0–100 μg/g of each element. Each standard was scanned over an area of 500 × 500 μm with a step size of 100 μm in both the X and Z directions. The count rate of fluorescence X-rays (counts s⁻¹) corresponding to the limit of quantification for each element was determined from the resulting calibration curve.

### Analysis of ferritin gene expression in radular tissue

We analyzed previously reported radular tissue RNA-seq datasets from *A. japonica*, *Acanthochitona achates*, *Placiphorella stimpsoni*, and *C. stelleri* ^7^. The sequencing data were obtained from the DNA Data Bank of Japan (DDBJ) under the accession numbers DRR598739, DRR598778, DRR598779, and DRA007106.

For the *A. japonica* dataset, sequencing errors in the raw reads were corrected using Rcorrector (ver. 1.0.4) ^59^ with default settings, and reads flagged as uncorrectable were removed using TranscriptomeAssemblyTools [https://github.com/harvardinformatics/TranscriptomeAssemblyTools]. Adapter sequences and low-quality bases (Phred score < 5) were removed using Trim Galore! (ver. 0.6.5). A recent taxonomic reassessment proposed that the species historically referred to as *Acanthopleura japonica* (currently accepted as *Liolophura japonica*) comprises three species: *L. japonica, L. koreana, and L. sinensis* ^60^. The reference genome originally published as *L. japonica* ^33^ was subsequently revised to *L. sinensis* ^61^. The cleaned reads were aligned to this reference genome (RefSeq accession number: GCF_032854445.1) ^33^ using STAR (ver. 2.7.11b) ^62^ within the RSEM pipeline (ver. 1.3.3) ^63^. Gene-level read counts and transcript abundance, expressed as transcripts per million (TPM), were estimated using RSEM. Read counts and mapping statistics are provided in Supplementary Table 3. For *A. achates*, *P. stimpsoni*, and *C. stelleri*, de novo transcriptome assemblies were constructed from the cleaned reads as described previously ^7^. The cleaned reads from each species were pseudoaligned to their respective transcriptome assemblies using Kallisto (ver. 0.43.1) ^64^, and transcript abundance was estimated as TPM.

Homology searches against the reference genome and the de novo transcriptome assemblies of the other three species were performed using TBLASTN implemented in BLAST+ (ver. 2.14.0) ^65^, with known ferritin amino acid sequences as queries. The query sequences were those used in a previous phylogenetic analysis of ferritins from *C. stelleri* ^21^.

### Phylogenetic and subcellular localization analyses of ferritin

Two separate phylogenetic analyses were performed. First, to classify the *A. japonica* ferritins in a broad evolutionary context, a phylogenetic tree was constructed using 53 ferritin amino acid sequences from *A. japonica*, chordates, and arthropods, with a poriferan ferritin sequence as the outgroup (Fig. 3a; Supplementary Data 1).

Second, to classify ferritins expressed in radular tissue and compare their expression among chiton species, a separate phylogenetic analysis was performed using ferritin sequences from *A. japonica*, *A. achates*, *P. stimpsoni*, and *C. stelleri* (Supplementary Fig. 1). For the sequences identified from the de novo–assembled transcriptomes of *A. achates*, *P. stimpsoni*, and *C. stelleri*, redundant sequences were clustered using CD-HIT (ver. 4.8.1) ^66^ with a sequence identity threshold of 95%. Sequences shorter than 150 amino acids were excluded from subsequent analyses. The amino acid sequences of the ferritins used for phylogenetic tree construction are listed in Supplementary Data 4.

For both analyses, amino acid sequences were aligned using MAFFT (ver. 7.525) ^67^, and the resulting alignment was trimmed using trimAl (ver. 1.5.0) ^68^ with the -gappyout option. A maximum-likelihood phylogenetic tree was inferred using IQ-TREE (ver. 2.4.0) ^69^. The best-fit amino acid substitution model, LG+G4, was selected automatically using ModelFinder. Branch support was assessed using 10,000 bootstrap replicates. The resulting tree was visualized using iTOL (ver. 7.4.2; https://itol.embl.de/).

The subcellular localization of the identified *A. japonica* ferritins was evaluated using three prediction tools. N-terminal signal peptides and their cleavage sites were predicted using SignalP 6.0 ^70^. TargetP 2.0 ^71^ was used in parallel to distinguish signal peptides from mitochondrial transit peptides. Subcellular localization was independently predicted using DeepLoc 2.1 ^72^ (Supplementary Data 3).

### Prediction of iron-responsive elements and ferritin protein structures

The transcript sequences corresponding to the 10 *A. japonica* ferritin genes identified by homology searches were retrieved from NCBI. Iron-responsive elements (IREs) were predicted using SIREs (ver. 3.0) ^73^. The accession numbers of the analyzed transcripts and the prediction results are provided in Supplementary Data 2. Ferritin transcripts predicted to contain an IRE stem–loop with a conserved cytosine bulge at position 8 (C8), a contact site for iron regulatory proteins (IRPs), were indicated in the phylogenetic tree (Fig. 3b). Three-dimensional structures of the ferritin proteins were predicted using AlphaFold 3 ^74^. In the structural representations, only α-helical regions comprising at least three amino acid residues, all with pLDDT scores greater than 90, were displayed.

### Purification of ferritin from radular tissue and hemolymph

Radular tissues freshly dissected from 40 *A. japonica* individuals were pooled and gently triturated by repeated pipetting in 1 mL of PBS to dissociate the epithelial cells. The resulting epithelial cell suspension was centrifuged at 16,600 × g for 1 min at 4°C. The epithelial cell pellet was snap-frozen in liquid nitrogen and stored at −20°C until use. The pellet was resuspended in 100 µL of lysis buffer containing 50 mM Tris-HCl (pH 8.0), 1% Triton X-100, 150 mM NaCl, 2 mM ethylenediaminetetraacetic acid (EDTA), 1 mM sodium orthovanadate, 1 mM NaF, 2 µg/mL aprotinin, and 1× cOmplete EDTA-free Protease Inhibitor Cocktail (Roche, Mannheim, Germany), and sonicated on ice for 2 min. The lysate was incubated at 75°C for 15 min. After the addition of 100 µL of PBS, the sample was mixed by pipetting and centrifuged at 10,000 × g for 10 min at 4°C. The resulting supernatant was ultracentrifuged at 100,000 × g for 30 min at 4°C, and the pellet was resuspended in PBS to a final volume of 100 µL. The resuspended sample was subjected to size-exclusion chromatography (SEC) using a Superdex 200 Increase 10/300 GL column (Cytiva, Marlborough, MA, USA) equilibrated with PBS. Peak fractions containing ferritin were pooled and concentrated using an Amicon Ultra-0.5 centrifugal filter unit with a 100-kDa molecular-weight cutoff (Merck Millipore, Darmstadt, Germany). Protein concentrations were determined using a BCA Protein Assay Kit (TaKaRa Bio, Shiga, Japan) according to the manufacturer’s instructions.

For the purification of hemolymph ferritin, hemolymph was collected from the gills of three *A. japonica* using a syringe and incubated at 75°C for 15 min. After heat treatment, the hemolymph was processed by centrifugation, ultracentrifugation, SEC, and centrifugal concentration as described above for radular ferritin, without the cell dissociation, lysis, or sonication steps.

Purified radular and hemolymph ferritin preparations were applied to copper grids and negatively stained with 2% (w/v) aqueous uranyl acetate. Samples were examined using a Hitachi H-7650 transmission electron microscope (Hitachi, Tokyo, Japan) operated at an accelerating voltage of 80 kV.

### Polyacrylamide gel electrophoresis

Native PAGE was performed using 7.5% polyacrylamide gels as described previously ^18^. Gels were stained for protein with Coomassie Brilliant Blue R-250 (CBB) or for iron with a freshly prepared 1:1 (v/v) mixture of 2% (w/v) potassium ferrocyanide and 0.232 M HCl. Blue native PAGE (BN-PAGE) was performed using EzRun BlueNative (ATTO, Tokyo, Japan) according to the manufacturer’s instructions. Tricine SDS-PAGE was performed as described previously ^75^. Proteins separated by Tricine SDS-PAGE were electrotransferred onto a polyvinylidene difluoride (PVDF) membrane (Merck Millipore) for N-terminal sequence analysis.

### RNA extraction and quantitative reverse transcription PCR

Radular tissue, midgut gland, gonad, and foot were dissected separately from three *A. japonica* individuals. For radular tissue, regions containing the radular epithelium were collected under a stereomicroscope. Each tissue was immediately frozen in liquid nitrogen and stored at −80°C until use. Total RNA was extracted using the Monarch Total RNA Miniprep Kit (New England Biolabs, Ipswich, MA, USA) with DNase I (New England Biolabs) treatment. The purified RNA was reverse-transcribed into cDNA using LunaScript RT SuperMix (New England Biolabs) according to the manufacturer’s instructions.

Quantitative reverse transcription PCR (RT-qPCR) was performed using an Mx3000P Real-Time PCR System (Agilent Technologies, Santa Clara, CA, USA) and Luna Universal qPCR Master Mix (New England Biolabs). Target-gene expression was normalized to that of the actin reference gene, and relative expression levels were calculated using the ΔΔCt method ^76^. The primer sequences used for RT-qPCR are provided in Supplementary Table 4.

### RNA-seq analysis of ferritin gene expression in the midgut gland and foot

Midgut gland and foot tissues were dissected from *A. japonica,* and total RNA was extracted as described above. RNA quality was assessed using an Agilent 2100 Bioanalyzer (Agilent Technologies, Santa Clara, CA, USA). RNA-seq libraries were prepared using the TruSeq Stranded mRNA Sample Preparation Kit (Illumina, San Diego, CA, USA) and sequenced on an Illumina NovaSeq 6000 platform to generate paired-end reads. Raw reads were processed using the same quality-control procedure as that described above for the radular RNA-seq dataset. The resulting quality-filtered reads were mapped to the *A. japonica* reference genome using the same STAR–RSEM pipeline as that used for the radular RNA-seq data, and gene-level read counts and transcripts per million (TPM) values were calculated. Read counts and mapping statistics are provided in Supplementary Table 3.

Raw read counts were normalized using the trimmed mean of M values (TMM) method ^77^ implemented in edgeR (ver. 4.8.2) ^78^ under R (ver. 4.5.2). For heatmap visualization, TMM-normalized counts per million (CPM) were natural log-transformed after addition of a pseudocount of 1. The heatmap was generated using the pheatmap package (ver. 1.0.13) ^79^.

### Proteomic analysis of purified ferritin complexes

Purified radular and hemolymph ferritin fractions were separated by Tricine SDS-PAGE and stained with CBB. Each lane contained 6 µg of protein. For radular ferritin, the single major band at approximately 25 kDa was excised. For hemolymph ferritin, the three bands observed at approximately 25 kDa were excised and analyzed separately. After CBB destaining and cysteine alkylation, proteins were subjected to in-gel digestion with trypsin, and the resulting peptides were extracted. The peptide samples were desalted using GL-Tip SDB (GL Sciences, Tokyo, Japan) according to the manufacturer’s instructions.

Peptides were analyzed in data-dependent acquisition mode by nano-liquid chromatography– tandem mass spectrometry (nanoLC–MS/MS) using a Vanquish Neo UHPLC system (Thermo Fisher Scientific, Waltham, MA, USA) coupled to an LCMS-9050 quadrupole time-of-flight mass spectrometer (Shimadzu Corporation, Kyoto, Japan). The acquired MS/MS data were converted to mzML format using msConvert in ProteoWizard (ver. 3.0.25148) ^80^. Database searches were performed using MaxQuant (ver. 2.6.8.0) ^81^ against the *A. japonica* reference protein database associated with RefSeq genome assembly GCF_032854445.1 ^33^. Trypsin/P was specified as the proteolytic enzyme, and up to two missed cleavages were permitted. Carbamidomethylation of cysteine was specified as a fixed modification. Oxidation of methionine, acetylation of lysine and the protein N terminus, and pyroglutamate formation from glutamine or glutamic acid were specified as variable modifications. The minimum peptide length was set to seven amino acids. The false discovery rate was set to 1% using a reversed-sequence decoy database, and the match-between-runs function was enabled.

Protein abundance was estimated using intensity-based absolute quantification (iBAQ), in which the summed peptide intensities for each protein are divided by the number of theoretically observable tryptic peptides ^82^. Entries identified as reverse-sequence matches, potential contaminants, or proteins identified only by a modification site were removed using Perseus (ver. 2.1.4.0) ^83^. Proteins lacking the quantitative values required for subsequent comparisons were also excluded. The proteins retained after filtering and their quantitative values are provided in Supplementary Data 5, 6.

The relative abundance of each ferritin subunit was calculated as a relative iBAQ (riBAQ) value by dividing its iBAQ value by the sum of the iBAQ values of all ferritin subunits detected in the same sample ^84^. For hemolymph ferritin, the iBAQ value of each subunit was first calculated by summing its values across the three gel bands before calculating the riBAQ value.

### Recombinant expression, purification, and cryo-EM structure determination of AjsFer7

The coding sequence of AjsFer7 was cloned into pET-47b (+) by In-Fusion cloning using the primers listed in Supplementary Data 7, and the resulting construct was verified by DNA sequencing. The plasmid was transformed into *Escherichia coli* BL21(DE3) (Novagen, Madison, WI, USA). Cells were grown at 37 °C in LB medium containing 50 μg mL^-1^ kanamycin to an OD₆₀₀ of 0.5–0.6, and protein expression was induced with 0.1 mM IPTG at 20 °C for 20 h. Cells harvested from a 1-L culture were resuspended in 50 mL of lysis buffer (10 mM sodium phosphate, 500 mM NaCl, pH 7.8) and disrupted by sonication on ice. The clarified lysate was filtered through a 0.45-μm membrane and applied to 2 mL of TALON metal-affinity resin. After washing with 30 column volumes of lysis buffer, bound protein was eluted with lysis buffer containing 10 mM imidazole. The eluate was adjusted to 150 mM NaCl with a buffer containing 50 mM sodium phosphate, 150 mM NaCl, and 10 mM reduced glutathione (pH 7.8), and applied to a Resource Q column (Cytiva). The flow-through was concentrated and purified by a Superdex 200 Increase 10/300 GL column in 50 mM sodium phosphate, 150 mM NaCl, and 10 mM reduced glutathione (pH 7.8). Purified AjsFer7 (3.0 μL; approximately 11 mg mL^-1^) was applied to a plasma-cleaned Quantifoil R1.2/1.3 300-mesh Au grid, blotted for 5 s at 18 °C and 100% humidity, and vitrified in liquid ethane using a Vitrobot Mark IV. Movies were acquired using a Talos Arctica G2 microscope operated at 200 kV with a Falcon 4 detector in electron-counting mode. Data were collected at 0.84 Å per pixel over a defocus range of −0.8 to −2.0 μm, with a total dose of 50 e^-^Å^-2^ fractionated into 44 frames.

A total of 4,100 movies were processed using RELION (ver. 5.0.0) ^85, 86^, with CTF estimation using CTFFIND (ver. 4.1.14) ^87^ and particle picking with crYOLO ^88^. After 2D and 3D classification, CTF refinement, and Bayesian polishing, 194,659 of 799,275 particles were used for final refinement. Refinement with octahedral symmetry and Blush regularization, followed by post-processing, yielded a 2.00-Å map (FSC 0.143). A protomer from an AlphaFold 3 ^74^-predicted AjsFer7 octamer was expanded into a 24-mer using octahedral symmetry and fitted into the map in ChimeraX ^89^. The model was adjusted in ISOLDE (ver.1.8) ^90^, refined in real space with NCS constraints and validated using PHENIX (ver. 1.21.2) ^91, 92^.

### Motif discovery and structural mapping

The seven secretory ferritin subunits identified in hemolymph by proteomic analysis were divided into two groups: four also detected in native radular ferritin complexes (radula-associated subunits) and three detected only in hemolymph (hemolymph-only subunits). Motif discovery was performed using MEME in discriminative mode ^93^, with the four radula-associated subunits used as the primary sequence set and the three hemolymph-only subunits used as the control set. MEME was used to identify the six highest-ranking motifs. The occurrence of these motifs in all seven secretory ferritin sequences was subsequently evaluated using FIMO ^93^. Motif matches with a *P* value below 10^-30^ were retained. Motifs detected in the radula-associated subunits but not in any of the hemolymph-only subunits were defined as candidate radula-associated motifs (Supplementary Data 8).

The three-dimensional structure of AjsFer3, a radula-associated ferritin subunit identified by N-terminal sequencing and proteomic analysis, was predicted using AlphaFold 3 ^74^. The locations of the candidate motifs were mapped onto the predicted AjsFer3 structure and visualized using UCSF ChimeraX (ver. 1.10.1) ^89^. To visualize the motifs in the context of the ferritin 24-mer assembly, the AlphaFold 3 model of AjsFer3 was superimposed onto the cryo-electron microscopy structure of recombinant AjsFer7 using UCSF ChimeraX. Amino acid residues predicted to participate in protein– protein interaction interfaces were identified using PeSTo ^34^.

### In vivo ferritin tracking

Hemolymph ferritin was purified from *A. japonica* as described above, except that the Superdex 200 Increase 10/300 GL column (Cytiva) was equilibrated with HEPES buffer. Purified hemolymph ferritin and horse spleen ferritin (Sigma-Aldrich, St. Louis, MO, USA) were independently labeled with Alexa Fluor 488 NHS ester (Thermo Fisher Scientific, Waltham, MA, USA). Briefly, 3 µL of Alexa Fluor 488 NHS ester solution (10 mg/mL; 30 µg of dye) was added to 200 µL of each ferritin solution (2 mg/mL; 0.4 mg of protein), and the labeling reaction was performed according to the manufacturer’s instructions. Unreacted dye was removed, and each labeled ferritin preparation was buffer-exchanged into Ca^2+^- and Mg^2+^-free artificial seawater (CMF-ASW) and adjusted to 2 mg/mL using an Amicon Ultra-0.5 centrifugal filter unit with a 100-kDa molecular-weight cutoff (Merck Millipore). CMF-ASW was prepared based on previously described molluscan tissue culture media ^94, 95^, with Ca²⁺ and Mg²⁺ omitted to minimize protein aggregation. CMF-ASW contained 520 mM NaCl, 10 mM KCl, and 20 mM HEPES in Milli-Q water and was adjusted to pH 7.8–7.9. Each *A. japonica* individual was injected with 50 µL of labeled ferritin solution, corresponding to 100 µg of protein. The solution was injected into the hemolymph through a gill located between the girdle and foot. Following injection, the chitons were maintained for 48 h at 18°C in seawater collected near their natural habitat.

### Localization analysis of fluorescently labeled ferritin

At 48 h after injection, radular tissues were dissected from *A. japonica*, washed with CMF-ASW, and immediately fixed in 4% paraformaldehyde in PBS for 6 h at 4°C. The fixed tissues were cryoprotected overnight in 30% (w/v) sucrose in PBS at 4°C. All procedures were performed under light-protected conditions. The tissues were embedded in optimal cutting temperature (O.C.T.) compound (Sakura Finetek, Tokyo, Japan), and 10-µm-thick cryosections were prepared using a Leica CM1100 cryostat (Leica Biosystems, Nussloch, Germany). Bright-field and fluorescence images were acquired using an Olympus IX83 inverted microscope and an FV3000 confocal laser scanning microscope (Olympus, Tokyo, Japan), respectively. Eight consecutive teeth, corresponding to teeth #−3 to #4 according to the numbering scheme defined above, were examined. Fluorescence intensity was quantified using Fiji ^58^ at the positions indicated in the corresponding images. To facilitate comparison among tooth positions, each fluorescence intensity value was divided by the maximum value measured across all analyzed images. Endogenous ferritin granules in the bright-field images were quantified using red chromaticity (“redness”), calculated as (R/(R+G+B)), where (R), (G), and (B) represent the intensities of the red, green, and blue channels, respectively.

### Preparation and injection of Ag-loaded hemolymph ferritin

Hemolymph ferritin was purified from *A. japonica* as described above, except that the SEC column was equilibrated with 20 mM Tris-HCl (pH 7.3) containing 50 mM NaCl ^96^. Iron was removed from the purified ferritin using a previously described method with modifications ^97^. The deironing solution was prepared by mixing 30 µL of 70 mM NaCl, 10 µL of 60% ammonium thioglycolate, and 200 µL of 0.3 M EDTA, followed by adjustment of the pH to 7.0. Purified hemolymph ferritin solution (125 µL) was added to the deironing solution and incubated for 3 h at 4°C.

Following deironing, the sample was buffer-exchanged into 20 mM HEPES (pH 7.2) to remove residual reducing and chelating agents and minimize the chloride concentration before Ag loading. Buffer exchange was performed using an Amicon Ultra-0.5 centrifugal filter unit with a 100-kDa molecular-weight cutoff (Merck Millipore) ^98^. Five buffer-exchange cycles were performed, each consisting of centrifugal concentration at 10,000 × g for 5 min at 4°C followed by redilution with 20 mM HEPES. The protein concentration of the recovered deironed ferritin was determined using the BCA assay and adjusted to 1.0 mg/mL in a final volume of 200 µL.

For Ag loading, 20 µL of 40 mM AgNO₃ was added to the deironed ferritin solution, and the mixture was incubated for 60 min at room temperature. Subsequently, 40 µL of 10 mM NaBH₄ was added, and the mixture was incubated for an additional 3 h at room temperature. These Ag-loading conditions were adapted from a previously described method with modifications ^98^. The reaction mixture was buffer-exchanged into 20 mM HEPES (pH 7.2) using the same five-cycle ultrafiltration procedure described above, and the final sample volume was adjusted to 200 µL. The post-loading purification procedure was adapted from previously described methods ^99, 100^.

Each *A. japonica* individual was injected with 50 µL of Ag-loaded hemolymph ferritin. The sample was injected into the hemolymph through a gill located between the girdle and foot. Following injection, the animals were maintained for 72 h at 18°C in seawater collected near their natural habitat. Radular tissues were then dissected, and paraffin sections for SR-XRF analysis were prepared as described above.

## Data and materials availability

RNA-seq data from foot and midgut gland have been deposited in the DDBJ Sequence Read Archive under accession numbers DRR1213921 and DRR1213922, respectively. The cryo-EM density map and atomic coordinates of recombinant AjsFer7 have been deposited in the Electron Microscopy Data Bank (EMDB) and the Protein Data Bank (PDB) under accession codes EMD-84134 and 46VT, respectively.

## Acknowledgements

We thank K. Saito of the Ushimado Marine Institute, Okayama University, for assistance with the collection of live chiton specimens and H. Fukuda of Okayama University for valuable insights into chiton internal anatomy. We are grateful to S. Kishimoto, E. Sanehira, and M. Kajino of the Central Research Laboratory, Okayama University, for technical assistance with the preparation of paraffin-embedded sections, and to M. Tsukano of the same facility for technical assistance with TEM analysis of ferritin. We thank T. Shiokawa and H. Tada of the Division of Instrumental Analysis for assistance with N-terminal amino acid sequencing, and Y. Ikeuchi of the Department of Genomics and Proteomics, Advanced Science Research Center, Okayama University, for assistance with liquid chromatography–tandem mass spectrometry analyses. Computations were partially performed on the NIG supercomputer at the ROIS National Institute of Genetics. The synchrotron radiation experiments were performed at the BL37XU in SPring-8 under the Priority Nanotechnology Support Program administered by the Japan Synchrotron Radiation Research Institute (Proposal numbers 2025A1457 and 2026A1181). We thank O. Sekizawa for his support during SR-XRF measurements. We are grateful to T. Takami and Y. Tsuchiya of the Institute of Plant Science and Resources, Okayama University for their valuable assistance with the ICP–MS analysis. This research was partially supported by Research Support Project for Life Science and Drug Discovery (Basis for Supporting Innovative Drug Discovery and Life Science Research (BINDS)) from AMED under Grant Number JP26ama121001 (support numbers 6582 and 4525). We thank M. Sugiyama of Kyoto University for providing access to experimental facilities through the BINDS support program.

## Author contributions

Conceptualization: R.O., M.N. Funding acquisition: R.O., M.N., T.S. Investigation: R.O., M.S., K.Oka., N.A.R.V., T.N., Y.F., T.S., K.Oko., K.Ob., D.K., A.S., S.H.-T., M.N. Supervision: M.N. Writing – original draft: R.O., M.N. Writing – review & editing: R.O., M.S., T.N., Y.F., T.S., K.Oko., K.Ob., D.K., A.S., S.H.-T., M.N.

