## Supplementary information for "Secretory ferritins mediate iron delivery during magnetite tooth formation in chitons"

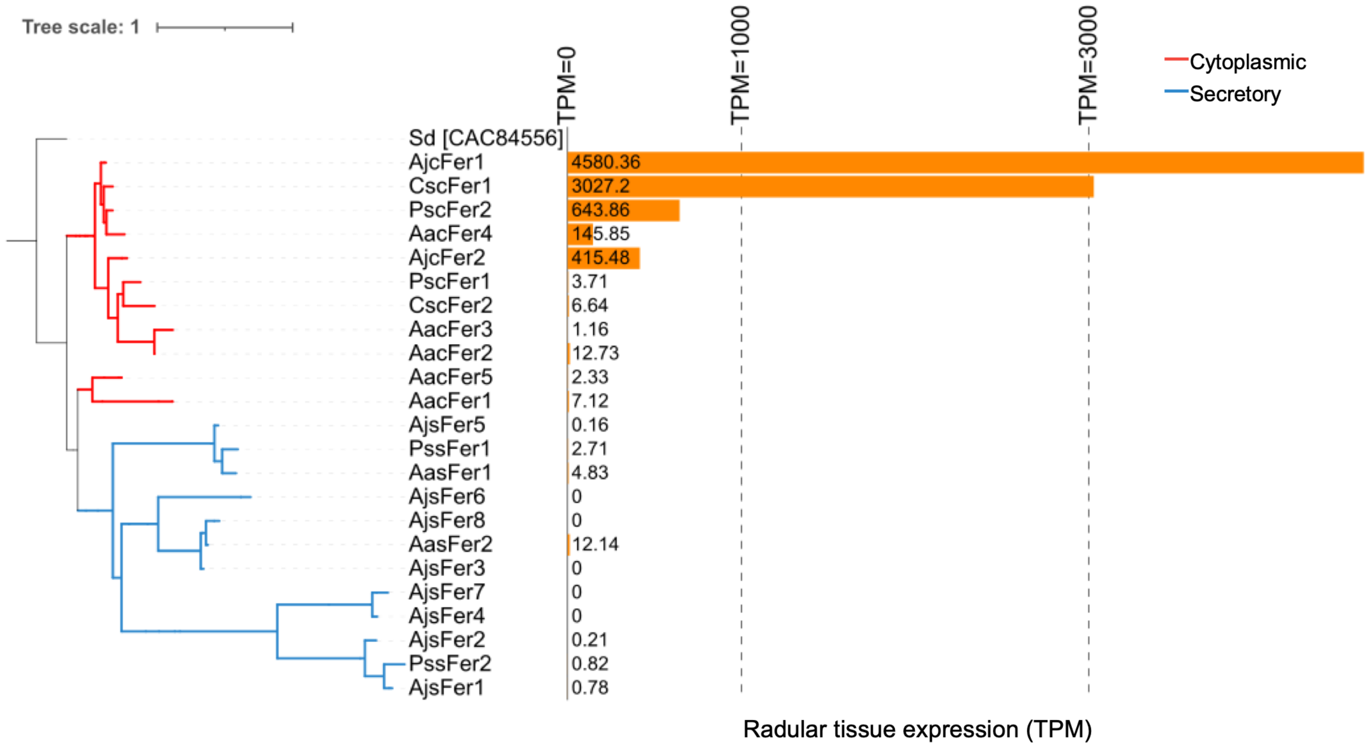

**Supplementary Fig. 1. Phylogenetic classification and expression levels (TPM) of ferritin genes in radular tissue of four chiton species based on transcriptome data.** The phylogenetic tree was constructed using the 10 *A. japonica* ferritin amino acid sequences shown in Fig. 3a together with ferritin amino acid sequences predicted from de novo-assembled transcripts of *Acanthochitona achates, Cryptochiton stelleri,* and *Placiphorella stimpsoni.* The corresponding transcript IDs are listed in Supplementary Data 4. For *A. japonica*, transcripts per million (TPM) values were calculated by mapping quality-filtered reads to the reference genome. For the other three species, TPM values were calculated by pseudoaligning quality-filtered reads to their respective de novo-assembled transcriptomes using kallisto.

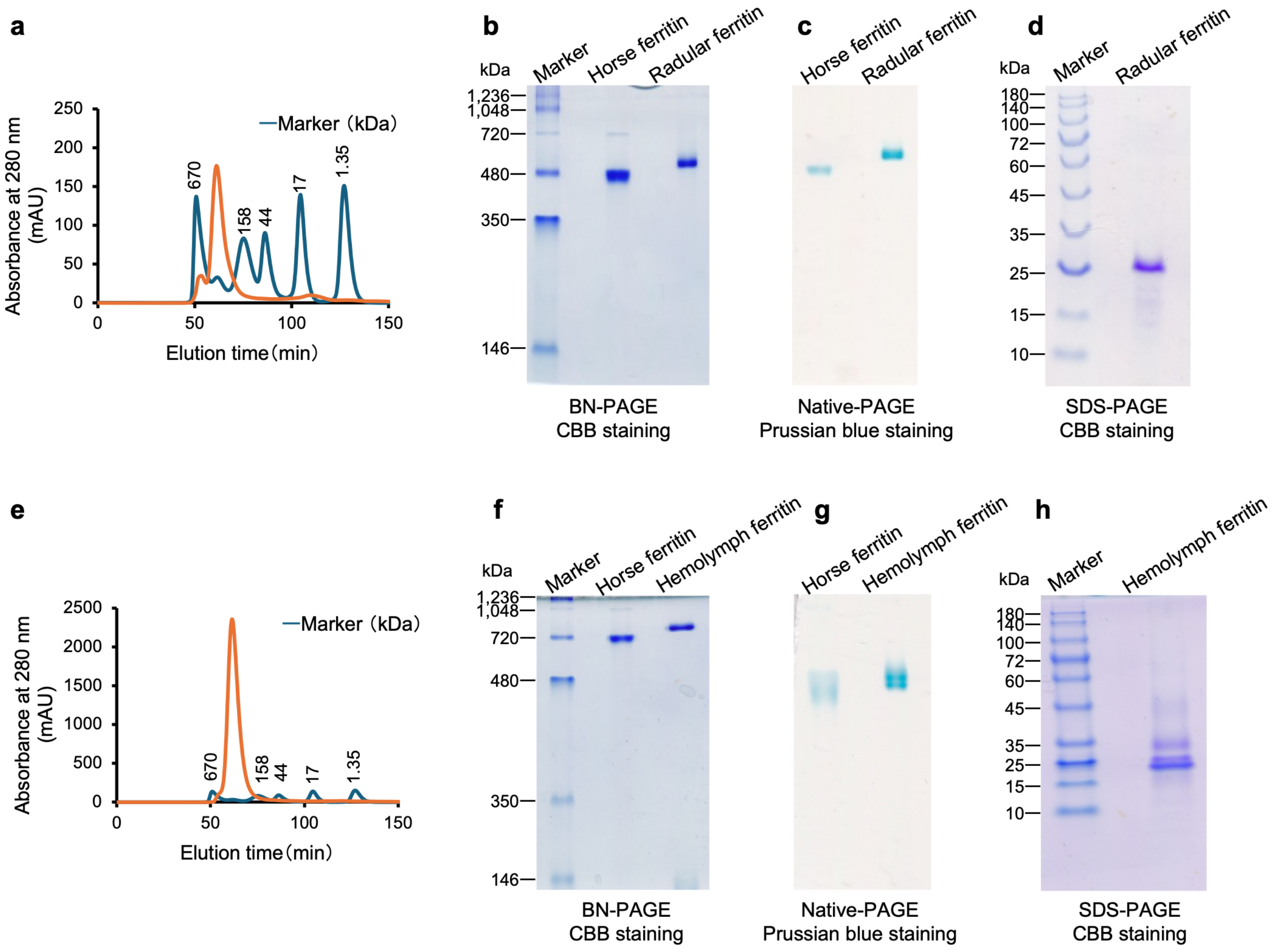

**Supplementary Fig. 2. Size-exclusion chromatography and PAGE analyses of ferritin isolated from radular tissue and hemolymph. a,** Size-exclusion chromatography elution profile of ferritin isolated from radular tissue. **b,** Blue native PAGE (BN-PAGE) with Coomassie Brilliant Blue (CBB) staining. **c,** Native PAGE with Prussian blue staining. **d,** Tricine SDS-PAGE with CBB staining. **e,** Size-exclusion chromatography elution profile of ferritin isolated from hemolymph. **f,** BN-PAGE with CBB staining. **g,** Native PAGE with Prussian blue staining. **h,** Tricine SDS-PAGE with CBB staining.

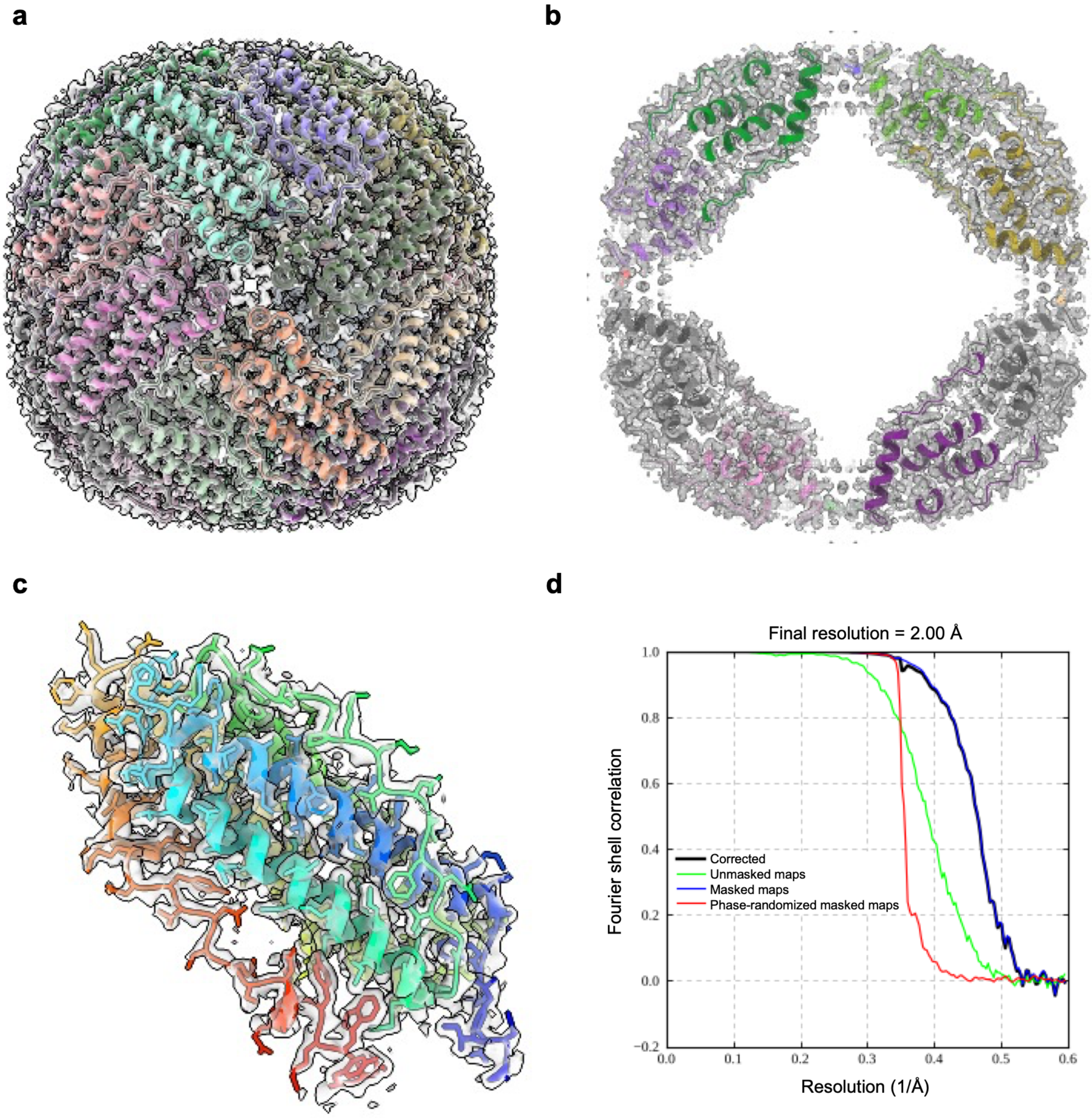

**Supplementary Fig. 3. Cryo-EM structural analysis of recombinant AjsFer7. a–c**, Cryo-EM map (EMD-84134; post-processed map contoured at 0.012) with the atomic model (PDB 46VT) superimposed. **a**, Overall view along a four-fold axis. The map is shown as a semi-transparent gray surface and the model as a cartoon colored by chain. **b**, Central slab viewed along a four-fold axis. The map is shown as a gray mesh and the model as a cartoon colored by chain. **c**, A single protomer (chain A), with the map shown within 2.5 Å of its atoms as a semi-transparent gray surface. Chain A is shown as sticks colored from the N terminus (blue) to the C terminus (red). **d**, Gold-standard Fourier shell correlation (FSC) curves from RELION post-processing for unmasked (green), masked (blue), phase-randomized masked (red; phases randomized beyond 2.89 Å) and masking-corrected (black) maps. The resolution of 2.00 Å was determined at FSC = 0.143.

**Supplementary Table 1.** Cryo-EM data collection, refinement and validation statistics.

| | AjsFer7<br>(EMD-84134)<br>(PDB 46VT) |
|---|---|
| **Data collection and processing** | |
| Magnification | 120,000 |
| Voltage (kV) | 200 |
| Electron exposure (e–/Å$^2$) | 50 |
| Defocus range (μm) | −0.8 to −2.0 |
| Pixel size (Å) | 0.84 |
| Symmetry imposed | O |
| Initial particle images (no.) | 799,275 |
| Final particle images (no.) | 194,659 |
| Map resolution (Å) | 2.00 |
| FSC threshold | 0.143 |
| **Refinement** | |
| Initial model used (PDB code) | AlphaFold 3 model |
| Model resolution (Å) | 2.0 |
| FSC threshold | 0.5 |
| Map sharpening *B* factor (Å$^2$) | −51.1 |
| Model composition | |
| Non-hydrogen atoms | 36,096 |
| Protein residues | 4,488 |
| Ligands | 0 |
| *B* factors (Å$^2$) | |
| Protein (min/max/mean) | 3.17/80.93/19.25 |
| R.m.s. deviations | |
| Bond lengths (Å) | 0.007 |
| Bond angles (°) | 0.889 |
| Validation | |
| MolProbity score | 1.07 |
| Clashscore | 2.46 |
| Poor rotamers (%) | 0.00 |
| Ramachandran plot | |
| Favored (%) | 97.84 |
| Allowed (%) | 2.16 |
| Disallowed (%) | 0.00 |

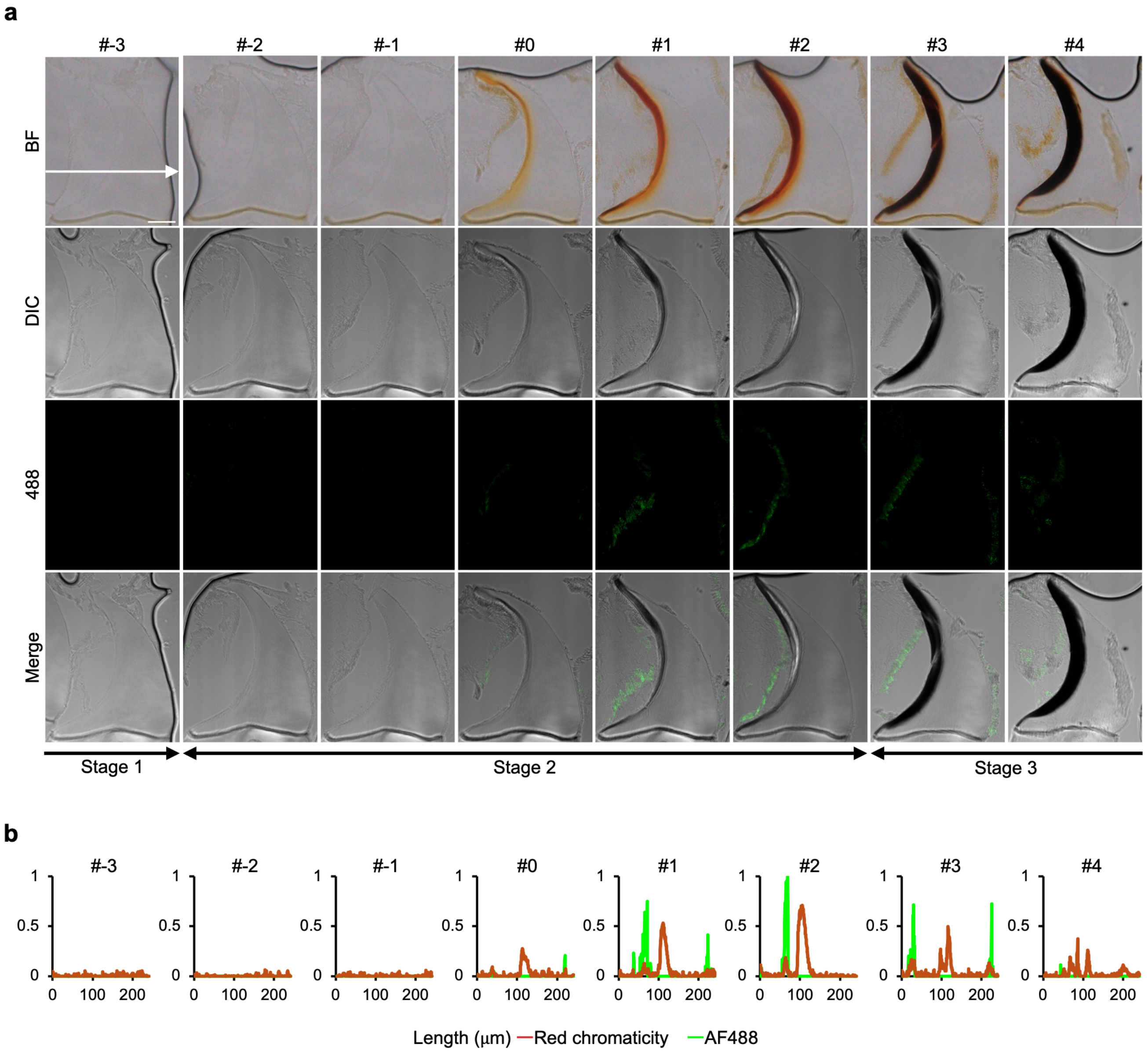

**Supplementary Fig. 4. In vivo tracking of fluorescently labeled chiton hemolymph ferritin in radular tissue. a,** Bright-field (BF) and fluorescence images of radular tissue sections from a chiton injected with Alexa Fluor 488-labeled hemolymph ferritin. Scale bar, 50 μm. **b,** Spatial profiles of Alexa Fluor 488 fluorescence (green), indicating the distribution of labeled ferritin, and red chromaticity (red), measured along the arrows shown in the corresponding BF images. Red chromaticity indicates the ferritin-rich granular regions surrounding the teeth and the iron oxide-rich regions within the teeth.

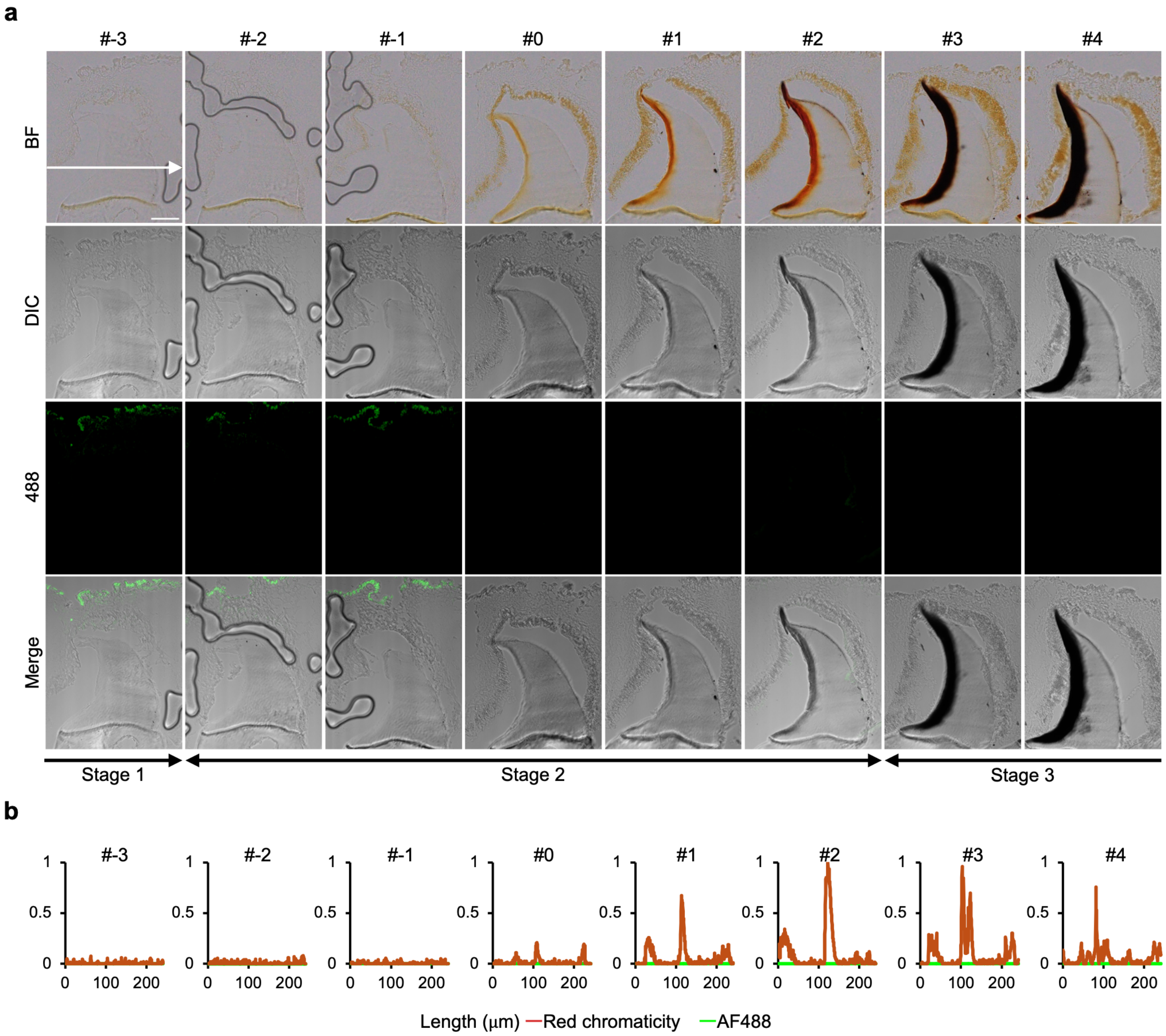

**Supplementary Fig. 5. In vivo tracking of fluorescently labeled horse spleen ferritin in radular tissue.** **a,** Bright-field (BF) and fluorescence images of radular tissue sections from a chiton injected with Alexa Fluor 488-labeled horse spleen ferritin. Scale bar, 50 μm. **b,** Spatial profiles of Alexa Fluor 488 fluorescence (green), indicating the distribution of labeled horse spleen ferritin, and red chromaticity (red), measured along the arrows shown in the corresponding BF images. Red chromaticity indicates the ferritin-rich granular regions surrounding the teeth and the iron oxide-rich regions within the teeth.

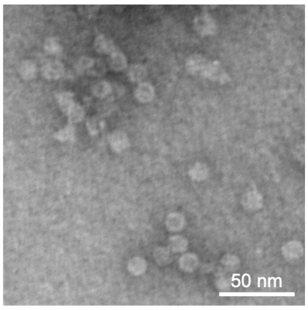

**Supplementary Fig. 6. TEM image of hemolymph ferritin after Ag loading.**

**Supplementary Table 2.** Iron and silver contents of native and Ag-loaded hemolymph ferritin determined by ICP-MS.

| Sample | Fe (ng/mg protein) | Ag (ng/mg protein) |
|---|---|---|
| Hemolymph Ferritin | 380.3 | 0.0 |
| Ag-loaded Hemolymph Ferritin | 26 | 85.9 |

**Supplementary Table 3.** RNA-seq read mapping statistics for the midgut gland, foot, and radular tissue of *Acanthopleura japonica.*

| Tissue | Midgut gland | Foot | Radula |
|---|---|---|---|
| Number of clean reads used for mapping | 38,145,849 | 38,168,350 | 41,799,908 |
| Uniquely mapped reads % | 83.16 | 79.45 | 71.16 |
| Reads mapped to multiple loci % | 12.35 | 12.24 | 22.66 |
| Overall mapping rate % | 95.51 | 91.69 | 93.82 |

**Supplementary Table 4.** Primer sequences used for RT-qPCR.

| Target gene | forward/reverse | Nucleotide sequences (5' to 3') |
|---|---|---|
| Actin gene | forward | AACCACAGCCGAGAGAGAAA |
| | reverse | ACCTGAATCTCTCGTTGCCA |
| AjsFer3（XM_064743856.1, proteinid=XP_064599926.1） | forward | CCATTGAGCACCATGTGAAC |
| | reverse | TATTCACCCAGACCGTCTCC |

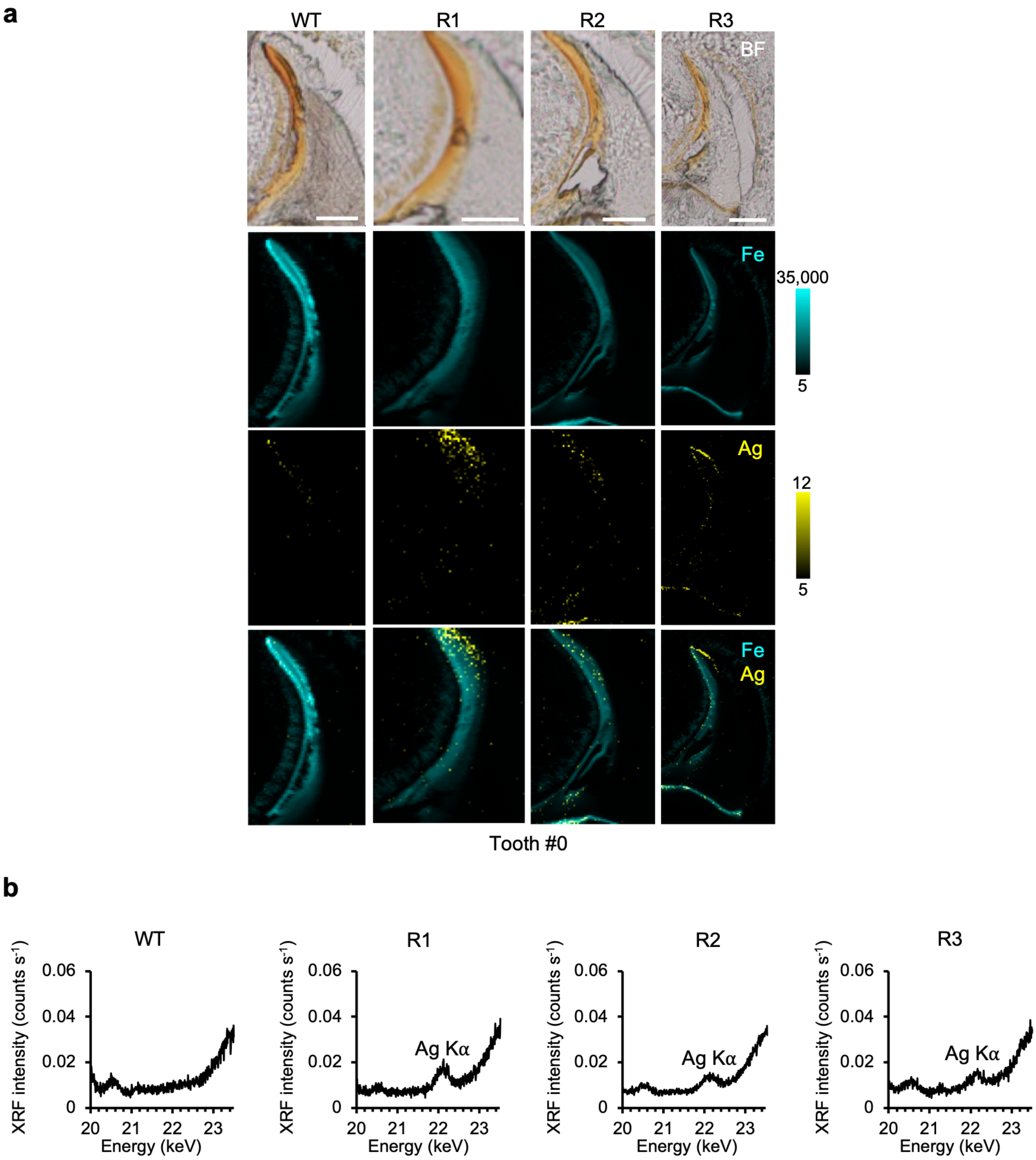

**Supplementary Fig. 7. SR-XRF elemental maps and XRF spectra of radular tissue from chitons injected with Ag-loaded hemolymph ferritin. a,** Ag and Fe maps of tooth #0 from an uninjected control (WT) and three individuals injected with Ag-loaded hemolymph ferritin (R1–R3). The color bars indicate the X-ray fluorescence count-rate range from the limit of quantification to the maximum for each element (counts $s^{-1}$). Scale bar, 50 µm. **b,** XRF spectra acquired from the analyzed regions in **a**. The expected position of the Ag Kα line is indicated. The XRF spectra were normalized to the total acquisition time.

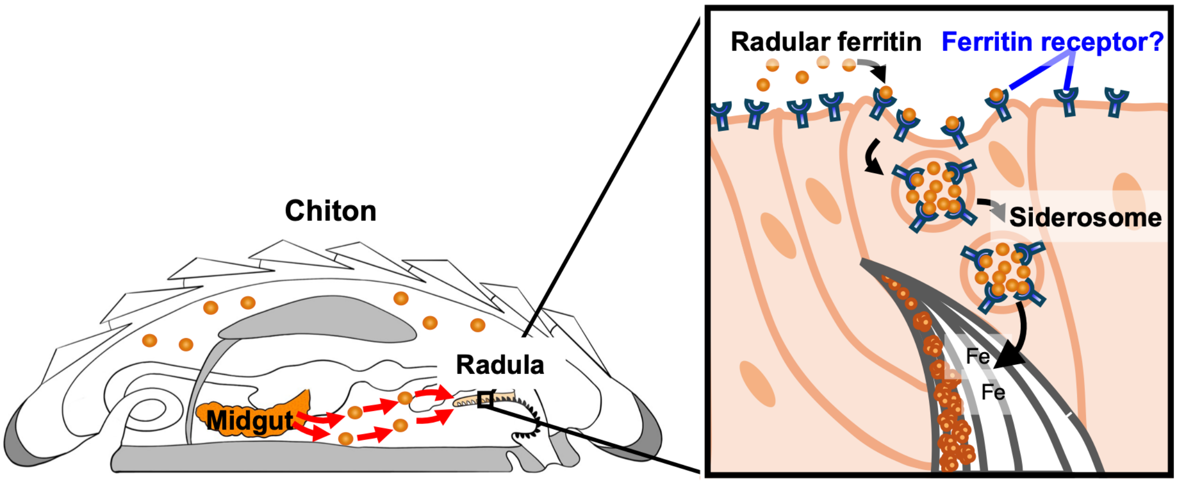

**Supplementary Fig. 8. Proposed model of secretory ferritin transport from the midgut gland to radular tissue via the hemolymph, based on the results of this study.**
